# Geometric deep learning on brain shape predicts sex and age

**DOI:** 10.1101/2020.06.29.177543

**Authors:** Pierre Besson, Todd Parrish, Aggelos K. Katsaggelos, S. Kathleen Bandt

## Abstract

The complex relationship between the shape and function of the human brain remains elusive despite extensive studies of cortical folding over many decades. The analysis of cortical gyrification presents an opportunity to advance our knowledge about this relationship, and better understand the etiology of a variety of pathologies involving diverse degrees of cortical folding abnormalities. Surface-based approaches have been shown to be particularly efficient in their ability to accurately describe the folded sheet topology of the cortical ribbon. However, the utility of these approaches has been blunted by their reliance on manually defined features in order to capture all relevant geometric properties of cortical folding. In this paper, we propose a deep-learning based method to analyze cortical folding patterns in a data-driven way that alleviates this reliance on manual feature definition. This method builds on the emerging field of geometric deep-learning and uses convolutional neural network architecture adapted to the surface representation of the cortical ribbon. MRI data from 6,410 healthy subjects obtained from 11 publicly available data repositories were used to predict age and sex via brain shape analysis. Ages ranged from 6-89 years. Both inner and outer cortical surfaces were extracted using Freesurfer and then registered into MNI space. Two gCNNs were trained, the first of which to predict subject’s self-identified sex, the second of which to predict subject’s age. Class Activation Maps (CAM) and Regression Activation Maps (RAM) were constructed to map the topographic distribution of the most influential brain regions involved in the decision process for each gCNN. Using this approach, the gCNN was able to predict a subject’s sex with an average accuracy of 87.99% and achieved a Person’s coefficient of correlation of 0.93 with an average absolute error 4.58 years when predicting a subject’s age.

## 1.1 Introduction

The complex relationship between the shape and function of the human brain remains elusive despite extensive studies of cortical folding over many decades. Although the exact mechanisms involved in the brain’s gyrification process are not completely understood, cortical folding is believed to be related to both the cytoarchitecture of the cortex as well as the tension gradient along developing axons. Gyrification begins at gestational week 16 when the fetal brain is experiencing rapid growth. It has been postulated that competing forces including rapid expansion of the cortical mantle, differential laminar growth within the cortical ribbon and tensions along rapidly elongating axons connecting remote brain areas play pivotal roles in this process (Caviness 1975, Rakic 1988, Van Essen 1997, Hilgetag and Barbas 2005, Kroenke and Bayly 2018). Given these contributing forces, gyrification is most certainly related to brain function although to what degree is undetermined (Fischl, Rajendran et al. 2008, Zilles and Amunts 2010). Beyond the obvious gyrification anomalies observed in developmental abnormalities including lissencephaly and polymicrogyria often associated with compromised neurologic function, far more subtle cortical folding variations have also been observed in epilepsy, autism, schizophrenia, bipolar disorder and major depressive disorder (Nordahl, Dierker et al. 2007, Besson, Andermann et al. 2008, Cachia, Paillère-Martinot et al. 2008, Penttilä, Paillère-Martinot et al. 2009, Schmitgen, Depping et al. 2019). It has been shown that cortical folding patterns in the early stages of embryologic gyrification are a predictor of later neurobehavioral functioning (Dubois, Benders et al. 2008). The relationship between cortical folding and brain function persists in the healthy adult and can be associated with temperament traits and reading abilities (Whittle, Allen et al. 2009, Cachia, Roell et al. 2018). Therefore, the analysis of cortical gyrification offers an opportunity to further investigate the dynamic relationship between brain structure and function over the life-span and to better understand the etiology of developmental and age-related pathologies.

Surface-based approaches have been shown to be particularly efficient in their ability to accurately describe the folded sheet topology of the cortical ribbon. Despite this, designing a method to extract all relevant geometric properties of cortical folding remains challenging. Previous methods have included local gyrification index (Schaer, Cuadra et al. 2008), cortical complexity (Thompson, Lee et al. 2005, Toro, Perron et al. 2008), fractal dimension (Im, Lee et al. 2006) and sulcus morphology (Kochunov, Mangin et al. 2005). Each provides quantitative measures of cortical folding, however, they are all constrained by manual design and suffer from an inability to model distant, potentially non-linear co-variations of cortical folding patterns. Spectral analysis of the cortical and sub-cortical surfaces overcomes some of these limitations but is limited by its inability to map findings in an anatomically informative way (Wachinger, Golland et al. 2015).

In this paper, we introduce a data-driven framework to extract the relevant geometric properties of cortical folding in order to determine the relationship between cortical folding and demographic features including age and sex. This method is an adaptation of the work of Defferrard et al. (Defferrard, Bresson et al. 2016), who provided an efficient implementation of convolution neural networks (CNNs) for graphs which has served as the foundation for the emerging field of geometric deep learning (Bronstein MM 2017). CNNs are a deep-learning network architecture which has been widely and successfully used in image, video and speech processing. One of the main strengths of CNNs is their ability to learn representations of the data with multiple levels of abstraction (LeCun, Bengio et al. 2015). However, traditional CNNs implicitly assume data is organized as a regular grid and therefore cannot be adapted when data lie on irregular or non-Euclidean domains such as when relationships between data points are represented by means of a graph. Graph convolutional neural networks (gCNNs) were introduced to overcome this limitation (Defferrard, Bresson et al. 2016), offering all of the same advantages of CNNs to non-Euclidean domains, therefore making gCNNs an ideal choice for surface-based analysis. In addition, gCNNs benefit from the rich selection of tools designed for traditional CNN applications including the identification and mapping of the nodes of the graph most involved in the decision process (Zhou, Khosla et al. 2016).

In this paper, we adapt gCNNs to brain MRI including data from several large cohorts of healthy subjects, acquired across multiple centers using different image acquisition parameters. The purposes of this study were to 1) examine the predictive power of gCNNs to predict age and sex using only cortical morphology from this large and diverse dataset and 2) to identify the most discriminative cortical regions involved in the network’s decision making.

## 2.1 Materials and methods

### 2.1.1 Subjects

MRI data were obtained from 11 publicly available data repositories and downloaded between November and December 2017. These datasets were selected based on the following criteria: 1) exclusive inclusion of healthy subjects, 2) availability of T1-weighted data for all subjects, 3) availability of basic demographic information including age (even if approximate) and sex for all subjects, 4) availability of data covering a large age range of subjects, and finally, 5) diversify of acquisition protocols including image quality and scanner manufacturers. Using these criteria to select publicly available data repositories, we identified the following datasets for inclusion: Autism Brain Imaging Data Exchange II (ABIDE II), Age-ility, Cambridge Centre for Ageing Neuroscience (CamCan), Consortium for Reliability and Reproducibility (CoRR), Dallas Lifespan Brain Study (DLBS), Brain Genomics Superstruct Project (GSP), Human Connectome Project (HCP), Information eXtraction from Images (IXI), MPI-Leipzig Mind Brain Body (MPI-LMBB), Enhanced Nathan Kline Institute – Rockland Sample (NKI-RS), and Southwest University Adult Lifespan Dataset (SALD). In order to avoid including duplicate data from a single subject only the baseline scans were selected from datasets that repeated imaging sessions (i.e., longitudinal or test-retest studies). Similarly, when datasets included data from multiple overlapping centers (ABIDE II and CoRR) care was taken to ensure no subjects were duplicated based on subject ID, imaging protocols, and descriptions provided on corresponding websites. Diversity of the acquisition parameters was intentionally sought as an important criterion in order to obtain a more broadly generalizable network. The goal of this work was to develop a tool that would be amenable to large-scale deployment and utilization, therefore, we actively sought heterogeneity within our data so as to not limit the resulting tool’s utilization to specific acquisition parameters (such as image resolution, noise or the type of acquisition) or to a specific scanner manufacturer or institution.

Data were aggregated, including T1-weighted image data as well as age and sex for each subject. Subject’s precise age was provided for most subjects however the GSP dataset (1570 subjects included) provided age data in 2-year bins and the MPI-LMBB dataset (318 subjects included) provided age data in 5-year bins. For these two datasets, subject’s age was set to the center of corresponding bins. Sources and descriptions for all datasets are provided in the supplemental material.

After data processing and quality assessment (see Data preparation), a total of 6410 subjects were included in the study and used for further analysis. The distribution of included subjects across different cohorts, as well as the distribution of age and sex are shown in Figure 1. The mean age was (mean ± SD) 33.32 ± 17.53 years and ranged from 6.7 to 89.1 years. Age distribution disproportionally included subjects between 18 – 35 years due to inclusion of the HCP and GSP cohorts. There were 3517 (54.9%) females and 2893 (45.1%) males in the final dataset.

**Figure 1.**
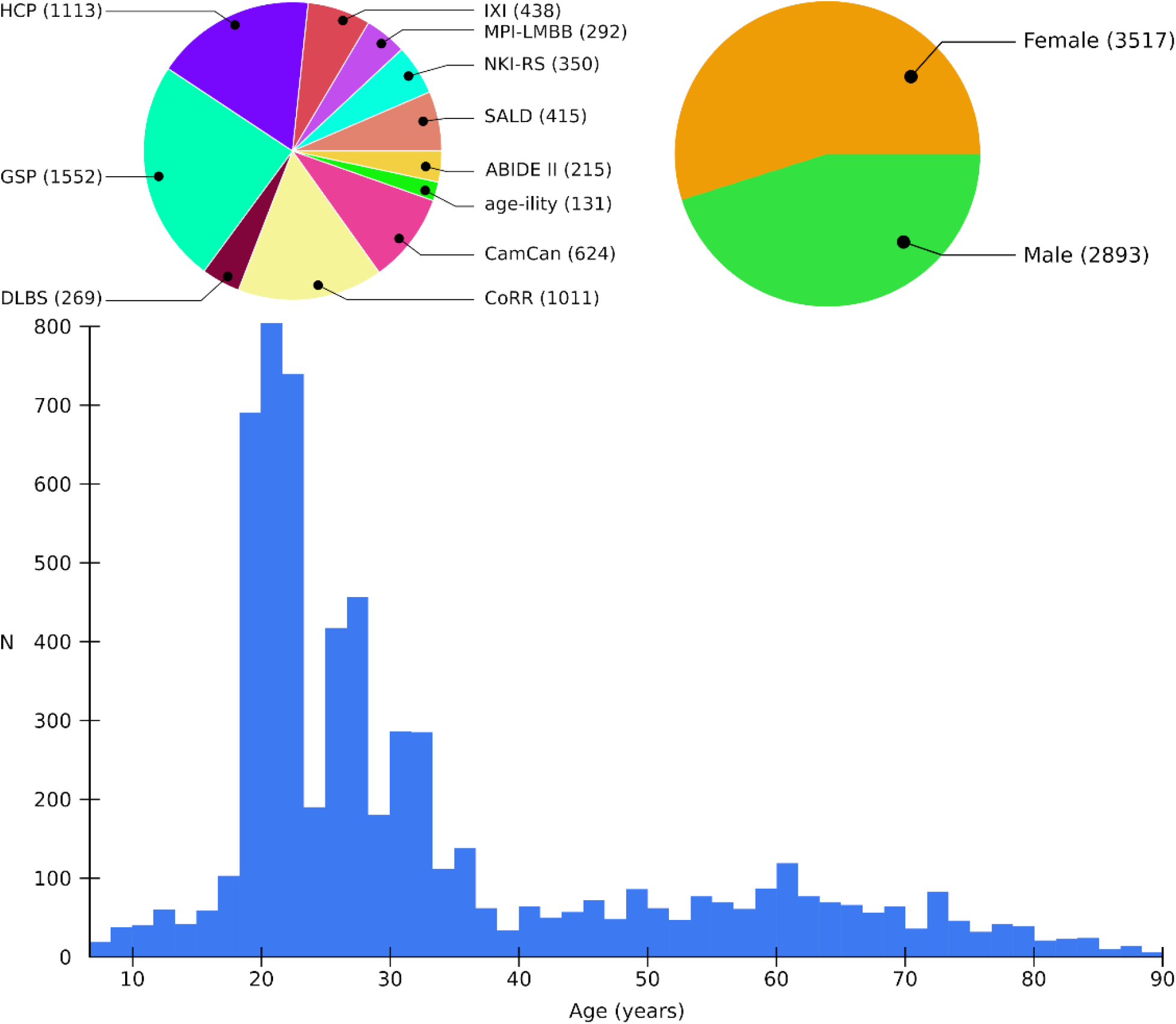
Composition of the final cohort included in the study according to the provenance of the dataset (N = 6410). The acronym of the dataset is indicated along with the number of included subjects from each dataset in parenthesis. Details about the datasets and MR images acquisition protocols are provided in supplemental data 1. In total, 3517 (58.9%) subjects were female. Histogram demonstrates age distribution of included subjects.

### 2.1.2 Data preparation

The overall data preparation process is illustrated in Figure 2. This included the following processing and quality assessment steps.

**Figure 2.**
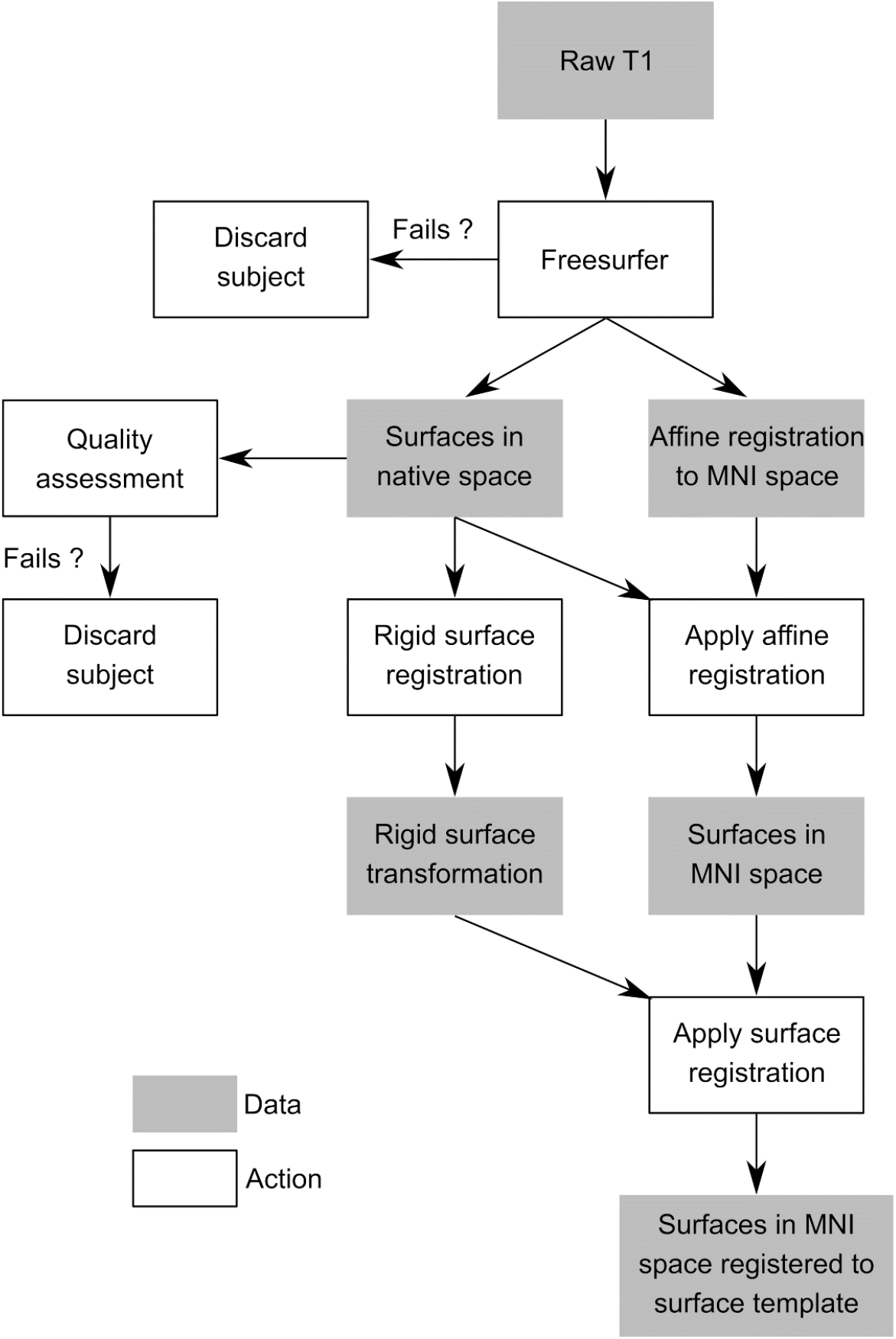
Flowchart of data preparation. After cortical surfaces are extracted using Freesurfer, they are affinely registered to the MNI space and rigidly registered with the surface template to ensure vertex-wise correspondence. Quality assessment ensures that the quality of surface extraction and registration is satisfactory.

#### 2.1.2.1 Image processing

T1 images were processed with Freesurfer (v6.0, https://surfer.nmr.mgh.harvard.edu) using Northwestern University’s High Performance Computing Cluster (QUEST, https://www.it.northwestern.edu/research/user-services/quest/) and the CBRAIN platform (Sherif, Rioux et al. 2014). Preprocessing steps included bias field correction, intensity normalization, spatial normalization, skull stripping and tissue segmentation (Dale, Fischl et al. 1999). The inner cortical surface, matching the white matter / grey matter junction, and the outer cortical surface, matching the grey matter / cerebro-spinal fluid interface, were then extracted. The surfaces were corrected for possible topological defects, inflated and parameterized (Fischl, Sereno et al. 1999, Fischl, Sereno et al. 1999).

#### 2.1.2.2 Quality assessment

Subjects were excluded from analysis if Freesurfer could not complete surface extraction on the first attempt. No attempt was made to reprocess subjects whose Freesurfer process failed before completion. For completed Freesurfer data, generated outputs were visually inspected to ensure that the cortical surfaces contained no obvious large errors. Subjects were excluded if at least one large error, significant enough to globally modify the sulcal pattern was identified. For example, subjects were discarded if one or both temporal poles were not entirely extracted, widespread inclusion of dura or if the surface was particularly noisy. Localized errors such as the inclusion of dura at the crown of a gyrus or local roughness were not disqualifying.

Finally, affine registrations to MNI space were inspected for all subjects. This was done by visually reviewing the cortical surfaces aligned with the MNI space. Incorrect registrations were manually corrected.

#### 2.1.2.3 Registration of the cortical surfaces to the MNI space

The four extracted cortical surfaces (left and right hemispheres, inner and outer cortical surfaces) were normalized by registering them to the MNI space (MNI305 template). This was done by applying the affine registration to the surface vertices coordinates to spatially normalize the cortical surfaces and normalize brain volumes.

#### 2.1.2.4 Surface registration

The purpose of surface registration is to establish a vertex-wise correspondence across individuals. By default, Freesurfer performs a non-rigid surface registration to improve cortical folding alignment (as determined by the curvature of the surfaces) between a subject and the surface template (Fischl, Sereno et al. 1999), which can result in distortion of the sulcal pattern. To preserve the initial sulcal folding pattern, surfaces were rigidly registered to the surface template (Freesurfer’s *fsaverage).* In a trade-off between precision of cortical alignment across subjects and preservation of individual subjects’ cortical folding pattern, this maneuver prioritized preservation of subjects’ folding pattern.

#### 2.1.2.5 Resampling surface coordinates to the common surface

Cartesian coordinates of the vertices (namely *X, Y* and *Z* coordinates of the inner and outer cortical surfaces) in individual’s native space were registered to the MNI space by applying the affine transformation, then they were registered to the common surface space using the rigid surface transformation. Finally, vertices coordinates were resampled to the freesurfer’s *fsaverage5* template to decrease the number of vertices in the common surface space to 10,242 vertices per hemisphere. This massive reduction in number of vertices was beneficial to train networks much faster while maintaining anatomic precision. Additionally, registration to the surface template permits mapping the cortical areas relevant in the decision process across subjects.

### 2.1.3 Training the Graph Convolutional Neural Network

Graph convolutional neural networks are a transposition of CNNs to any graph *G*. Let *G* = (*V, E, W*) a connected graph where *V* is a set of *n* vertices, *E* a set of edges and *W* an ℝ^*n×n*^ adjacency matrix such as *W_i,j_* > 0 if there is an edge between the nodes *i* and *j*, and *W_i,j_* = 0 otherwise. Let *F* ∈ ℝ^*n×m*^ be an input signal to the graph, so that as the *i*-th row of *F* is an m-dimensional vector assigned to the *i*-th node of *G*. The purpose of gCNNs is to process an input signal *F* mapped on a graph *G*.

The normalized graph Laplacian of *G* is defined as *L = I_n_* — *D*^−1/2^*WD*^−1/2^ where *I_n_* is the identity matrix of size *n × n* and *D* the diagonal degree matrix such as *D_i,i_* = ∑_*j*_ *W_i,j_*. The graph Laplacian is a real and symmetric matrix, therefore has a complete set of orthonormal eigenvectors ***u**_l_*, associated with eigenvalues *λ_l_*, such as *L**u**_l_* = *λ_l_**u**_l_*. The eigenvectors are known to be the graph Fourier modes and the eigenvalues the frequency spectrum (Hammond, Vandergheynst et al. 2011, Shuman, Narang et al. 2013). This enables the formulation of the Fourier transform 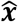 of a vector ***x*** mapped on the graph *G* as 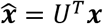, and its inverse 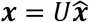, where *U* = [***u***_0_,…, ***u***_*n*-1_]. The convolution operator * can therefore be defined in the Fourier domain for two vectors ***x*** and ***y*** mapped on *G* such as 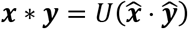, where · is the element-wise Hadamard product. As in Euclidean spaces, a filter can be expressed in the Fourier domain by a function *g* modulating the frequency spectrum to define ***y***, the vector obtained by filtering ***x*** with *g*, by: ***y*** = *Ug*(Λ)*U^T^**x*** with Λ = diag([λ_0_, …, λ_*n*-1_]). An efficient implementation of spectral filters *g* was proposed in (Hammond, Vandergheynst et al. 2011) which are computed recursively from the Laplacian and avoids its costly eigen decomposition. In addition, these filters are *K*-order polynomial parametrization, therefore requiring *K* parameters to define, and localized with support size *K*.

#### 2.1.3.1 Definition of the graph and input features

The graph provides the underlying structure of the data, in particular the relationships between its nodes, which is common to all subjects by construction. The cortical surfaces are triangulated meshes composed of 10,242 vertices per hemisphere connected to form triangles (see Figure 3). The nodes of the graph *G* were defined as the vertices of the cortical surfaces and the edges were the links of the triangles, weighted as a function of the length of the link such as 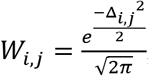, where Δ_*i,j*_ is the Euclidean distance between vertices *i* and *j*. The function *F,* defined at the nodes of *G* and constituting the input of the gCNN, were the vertices coordinates register to the MNI space, registered and resampled to the common surface space. Therefore, each vertex had 6 features if both the inner and outer cortical surfaces were used (*X, Y* and *Z* coordinates of both white and pial surfaces), or 3 features if only the inner cortical surfaces were used (*X, Y, Z* coordinates of the white surface only).

**Figure 3.**
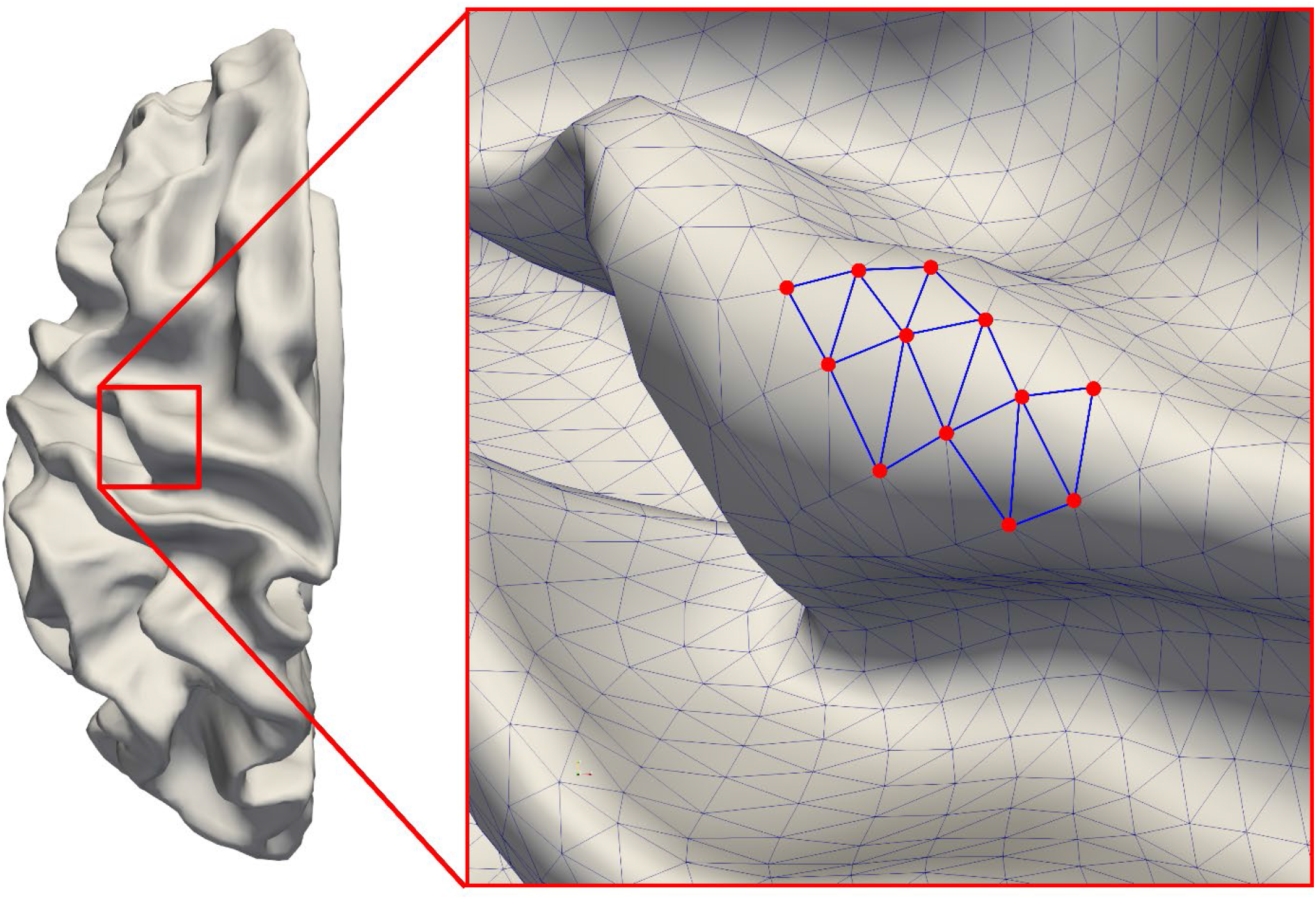
View of the left hemisphere surface template (freesurfer’s fsaverage5) and close up on a small part of the surface. Spatial relationships between each surface vertex are modeled via a graph. Nodes of the graph are surface vertices (red dots) and edges of the graph are the links between the vertices (blue lines). Edges are weighted as a function of their length. Node features, used as input of the gCNN, are each node’s Cartesian coordinates in MNI space, resampled to the common surface space. Therefore, the dimension of input features is either 6 when both the inner and outer cortical surfaces are used, or 3 when only the inner cortical surfaces are used.

#### 2.1.3.2 Graph pooling

The pooling operation intends to downsample a graph and summarize information within a neighborhood, therefore providing a multi-scale description of the data. This step was done as defined in (Defferrard, Bresson et al. 2016). In brief, setting the initial graph *G* and corresponding Laplacian matrix *L* to the level 0, multi-level graphs were obtained by a successive node clustering, so that a coarse graph was obtained by grouping pairs of nodes of its parent graph. After *B* successive clustering, this results in a set of graphs *G*_0_, *G*_1_,…, *G_B_* with associated Laplacian matrices *L*_0_, *L*_1_,…, *L_N_* such as *n_k_* = 2*n*_*k*-1_ where *n_k_* is the number of nodes of *G_k_*. The efficiency of the pooling operation is further increased by ordering the nodes so that node *p_i_* at level *k* + 1 has the two parent nodes *p*_2*i*_ and *p*_2*i*+1_ at level *k*.

#### 2.1.3.3 Architecture of the network

The design of our network architecture is shown in Figure 4. This novel gCNN architecture is inspired from the popular ResNet architecture which explicitly refers to the input to make the overall network easier to optimize (He, Zhang et al. 2016). This architecture A first batch normalization – graph convolution sequence extends the dimension of the input to the number of filters *F_c_*. Then a series of *B* blocks are applied, each of them reducing the number of nodes with a pooling operation. This is followed by a batch normalization, graph convolution with *F_L_* filters, and ReLU. Global average pooling (GAP) was used to summarize the *F_L_* feature maps into an *F_L_*-length vector, linearly combined with the final fully connected (FC) layer. The architecture of the blocks is illustrated in Figure 5.

**Figure 4.**
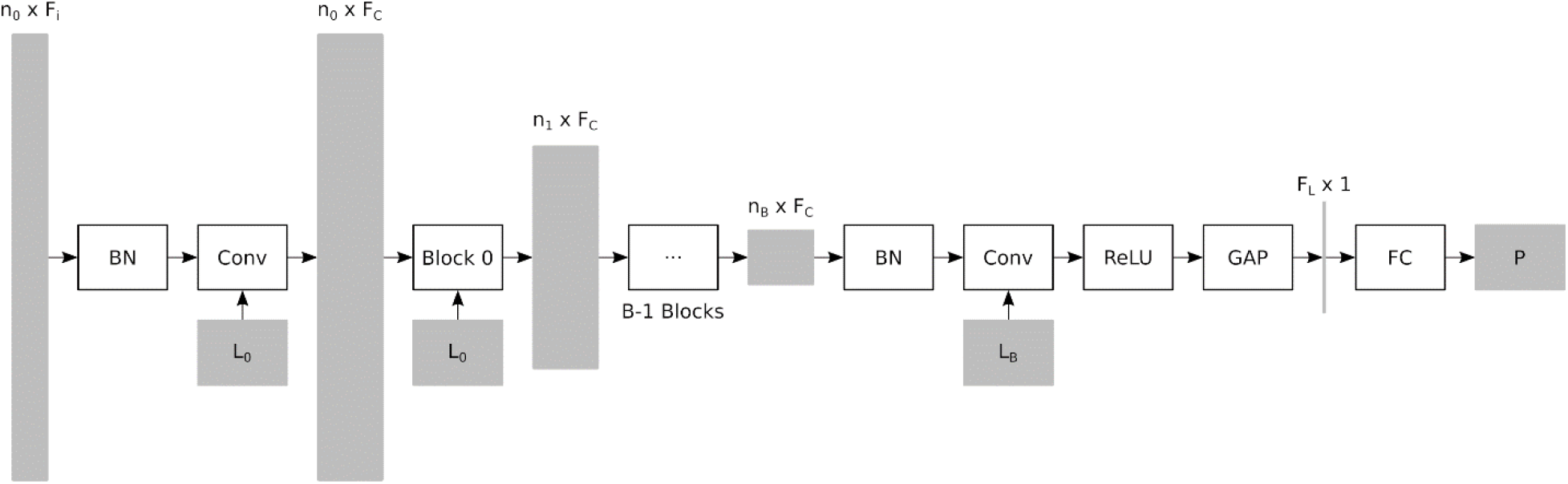
Architecture of the network. The input is of size *n*_0_ × *F_i_*, where *n*_0_ is the number of nodes of the graph at level 0 and *F_i_* the number of input features (either 6 or 3). Input is batch normalized (BN) and filtered using the Laplacian at level 0 (*L*_0_) and the graph convolution method (Conv). The purpose of the first convolution is to expend the number of features per node to *F_C_*. Then, *B* blocks are applied and for each block a pooling operation reduces the number of nodes such as *n_k_* = *n*_*k*-1_/2. After the last block, a graph convolution is applied with *F_L_* filters, followed by a rectified linear unit (ReLU) and a global average pooling (GAP) to obtain a feature vector of length *F_L_*. A linear combination of the elements of the feature vector is done by the fully connected layer (FC) and generates the prediction *P*.

**Figure 5.**
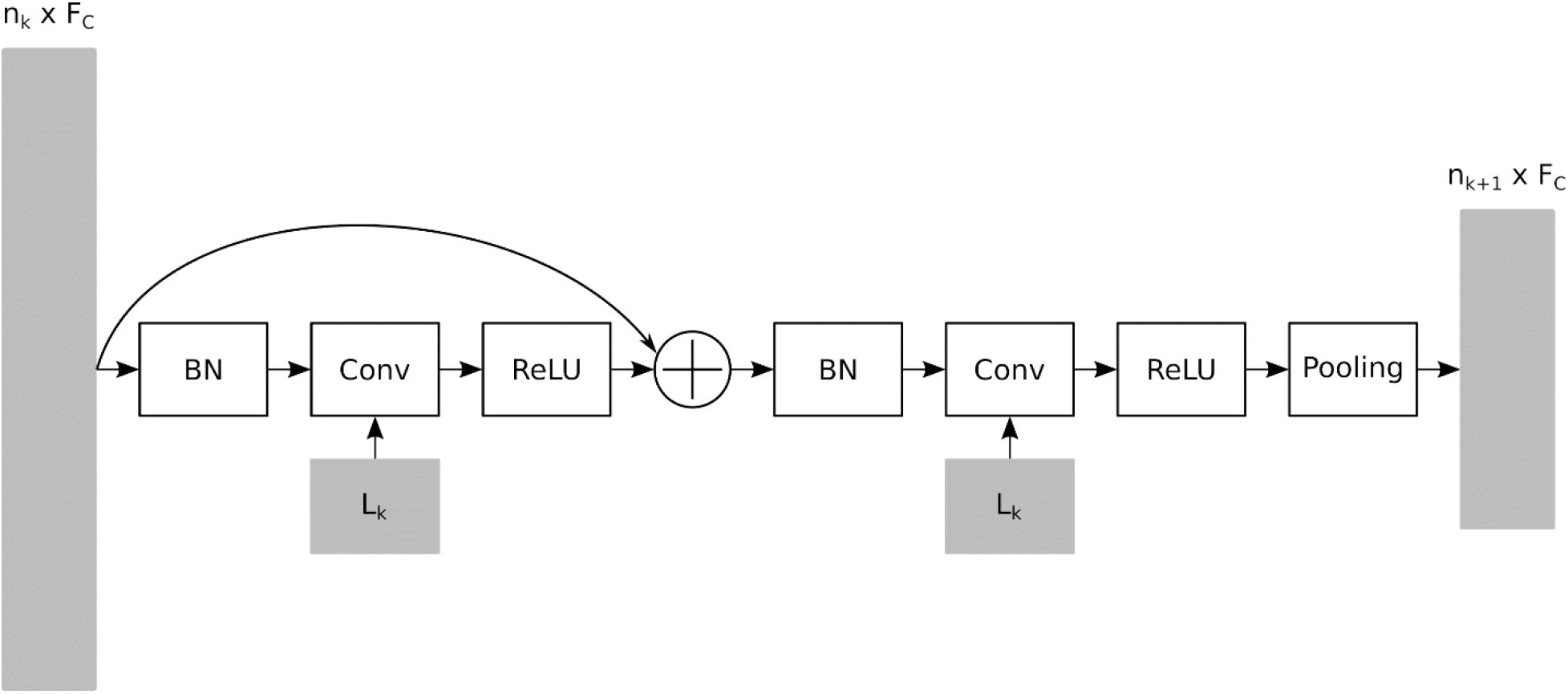
Architecture of a block. The input is added to the result of the first BN-Conv-ReLU operation. One more BN-Conv-ReLU sequence is followed by an average pooling, reducing the number of nodes to *n*_*k*+1_.

#### 2.1.3.4 Mapping the most influential brain regions

The topographic distribution of the most influential brain regions involved in the decision process are mapped using the Class Activation Maps (CAM) and Regression Activation Maps (RAM) approaches (Zhou, Khosla et al. 2016, Wang and Yang 2017). These maps are computed by interpolating the *F_L_*-length feature vectors obtained before GAP to the initial vector length *n*_0_ and creating a weighted sum of these vectors according to the weights of the FC layer. These maps qualitatively highlight the brain regions involved in the gCNN’s decision process.

### 2.1.4 Experiments

To evaluate the ability of gCNNs to learn meaningful features using only surface meshes of the cortex, both a classification and regression task were conducted. In both we fed the gCNN with either all four cortical surfaces including the inner and outer cortical surfaces from the left and right cerebral hemispheres leading to 6 input features per node, or only the inner cortical surfaces from each hemisphere which provide 3 features per node. Importantly, the use of the inner cortical surface only prevents the network from deriving any information, even indirectly, from the cortical thickness but instead forces the network to focus on morphologic descriptors of the cortical folding alone. In the case both the inner and outer cortical surfaces were used, the network can still extract morphologic descriptors of cortical folding, and is also given the possibility to extract descriptors of the inner cortical surfaces relative to the outer cortical surface, such as cortical thickness. The decision to use both of these training approaches was to determine the unique predictive value of cortical folding alone compared to richer information which may permit the network to infer additional measures of cortical morphology including cortical thickness.

#### 2.1.4.1 Sex prediction

As a first experiment, the network was trained to classify sex. Training and validation were handled via a L-fold cross-validation approach with *k* = 5, therefore the network was trained with 5128 instances and validated with 1282 independent instances.

For this task, the loss function was cross-entropy, *F_C_* was set to 32 filters, the number of blocks *B* set to 4 and *F_l_* was set to 128. In addition, the polynomial order was set to 5 for all filters, the learning rate was set to 0.001 with an exponential decay 0.95 every 400 iterations, filter weights were *L2* regularized to 5 × 10^−4^, dropout 0.5 was applied to the fully connected layer, the batch size was 64 and the optimization was performed with ADAM procedure (Kingma and Ba 2014).

#### 2.1.4.2 Age prediction

As a second task, the network was trained for age prediction using the same L-fold approach. The loss function was the mean squared error, *F_C_* was set to 32 filters, the number of blocks *B* was 7, *F_L_* was 16 and the polynomial order was 4 for all filters. The initial learning rate was 0.001 with an exponential decay 0.98 every 400 iterations, filter weights L2 regularization set to 1 × 10^−5^, dropout 0.5 was applied to the FC layer, the batch size was set to 64 and optimization was done with the ADAM method.

## 3.1 Results

### 3.1.1 Sex prediction

#### 3.1.1.1 Accuracy

Figure 6 shows the results obtained for each 5 folds using either all four cortical surfaces (inner and outer cortical surfaces for both the right and left cerebral hemispheres) or the bilateral inner cortical surfaces only. Using all cortical surfaces, the gCNN was able to predict a subject’s sex (using a binary classification scheme) with an average accuracy of 87.99%. Using only the bilateral inner cortical surfaces, average accuracy decreased only minimally to 85.23%. Table 1 reports the accuracy for sex prediction per dataset.

**Figure 6.**
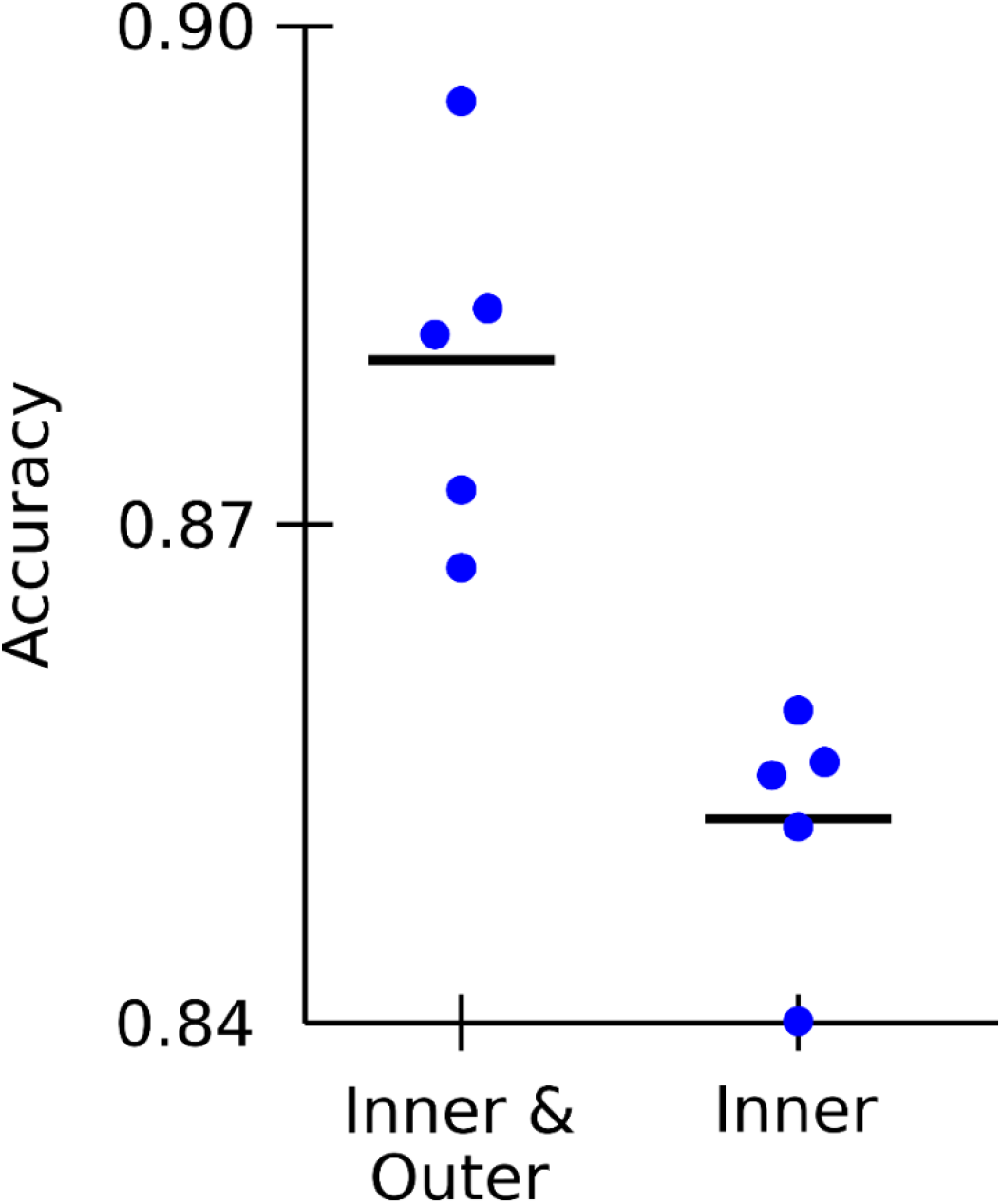
Results of the k-fold cross validation for sex classification, using either all cortical surfaces (inner and outer) or only the inner cortical surfaces. Bold lines indicate accuracy averages, which is 87.99% when all surfaces are used and 85.23% if only inner cortical surfaces are used.

#### 3.1.1.2 Mapping discriminative brain regions

Figure 7 shows the CAMs averaged across correctly classified subjects for male and female. Overall, there was a modest amount of overlap between brain regions predicting binary sex classification when using the bilateral inner cortical surfaces compared to classification when using the bilateral inner and outer cortical surfaces.

**Figure 7.**
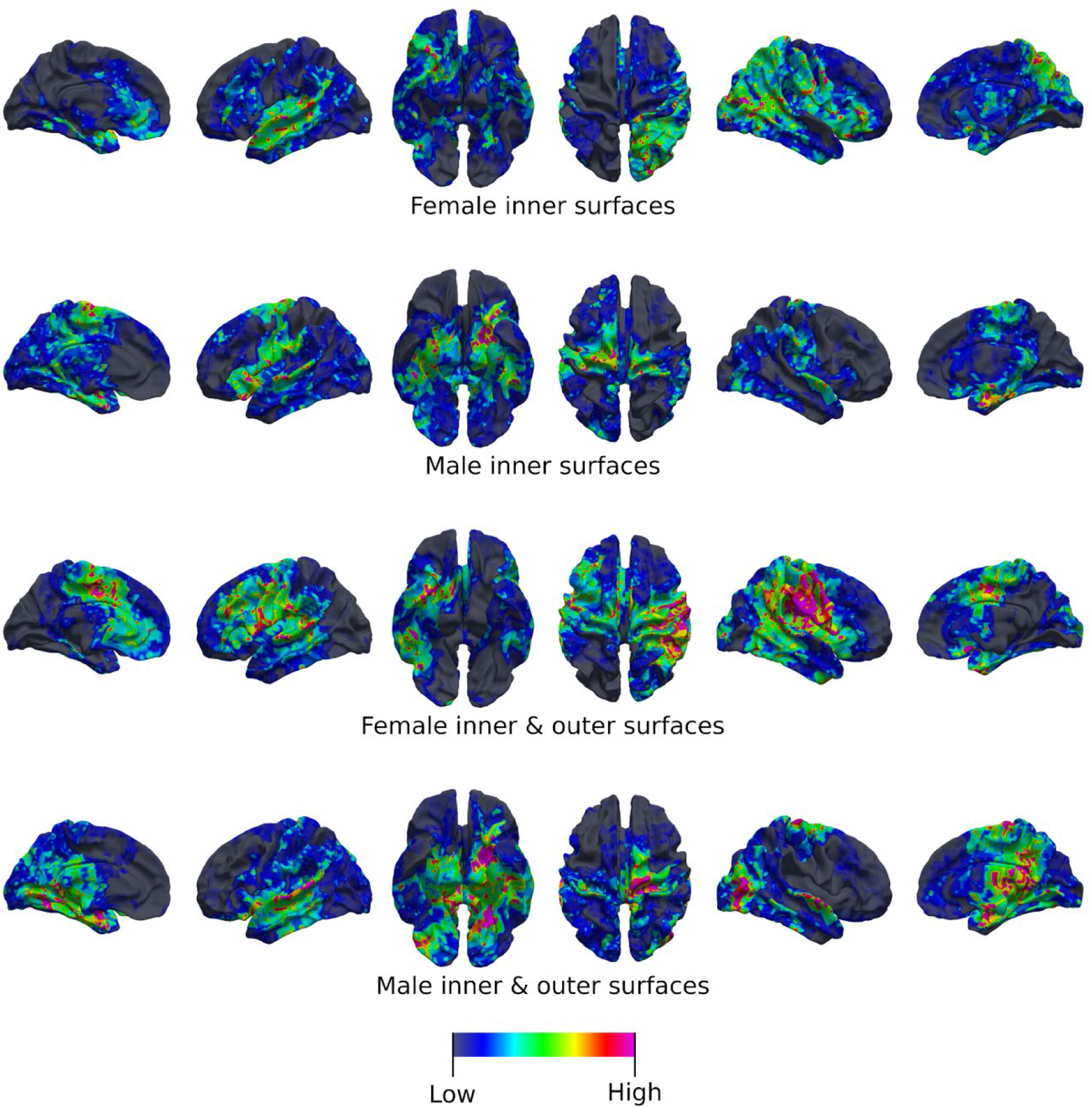
Average class activation maps (CAM) for correctly classified individuals using either only the inner cortical surfaces, or the inner and outer cortical surfaces. The color indicates cortical areas with either low or high involvement in the decision.

#### 3.1.1.3 Female classification

When using the bilateral inner cortical surfaces only (i.e., only using morphologic features of the brain’s sulcal/gyral folding pattern), the following brain regions were identified as predictive of female: left anterior cingulate cortex, left superior temporal gyrus, posterior left insula, right orbitofrontal cortex, entire right parietal cortex, right frontal operculum (including pars frontalis, pars triangularis and pars orbitalis) and the right insula.

When using the bilateral inner and outer cortical surfaces (i.e., morphologic features of the sulcal/gyral folding pattern and information regarding cortical thickness), the following brain regions were identified as predictive of female: left middle and anterior cingulate cortex, left superior temporal gyrus, left frontal operculum (including pars frontalis, pars triangularis and pars orbitalis), left posterior insula, right orbitofrontal cortex, right frontal operculum pars frontalis, right mid cingulate cortex, entire right parietal cortex and the right posterior insula.

#### 3.1.1.4 Male classification

When using the bilateral inner cortical surfaces only (i.e., only using morphologic features of the brain’s sulcal/gyral folding pattern), the following brain regions were identified as predictive of male: left medial paracentral lobule, left parahippocampal gyrus, left inferior frontal gyrus, left orbitofrontal cortex, left fusiform gyrus, left anterior insula, right medial paracentral lobule and the right medial temporal tip.

When using the bilateral inner and outer cortical surfaces (i.e., morphologic features of the sulcal/gyral folding pattern and information regarding cortical thickness), the following brain regions were identified as predictive of male: left medial temporal tip, left medial temporo-occipital junction, left superior temporal gyrus, left orbitofrontal cortex, left anterior insula, right medial paracentral lobule, right superior temporal gyrus, right lateral temporo-occipital junction and the right posterior cingulate cortex.

### 3.1.2 Age prediction

#### 3.1.2.1 Accuracy

Age prediction results for all 5 folds are illustrated in Figure 8. Using only the inner bilateral cortical surfaces, Pearson’s coefficient of correlation between the actual and predicted ages was 0.92 and the average absolute error was 4.91 years. Using all four cortical surfaces (bilateral inner and outer cortical surfaces), Person’s coefficient of correlation was 0.93 and the average absolute error 4.58 years. It is important to note that a small number of outliers were discarded from regression analysis. Table 2 reports the mean absolute error obtained for each dataset. Specifically, three subjects (0.05%) were excluded from regression analysis using only the bilateral inner cortical surfaces and six subjects (0.1%) were excluded from regression analysis using the bilateral inner and outer cortical surfaces due to their age prediction being off by 50 or more years. When the MRI data for the outliers were individually examined, nothing particularly abnormal was identified regarding their brain structure overall or their registration to MNI space.

**Figure 8.**
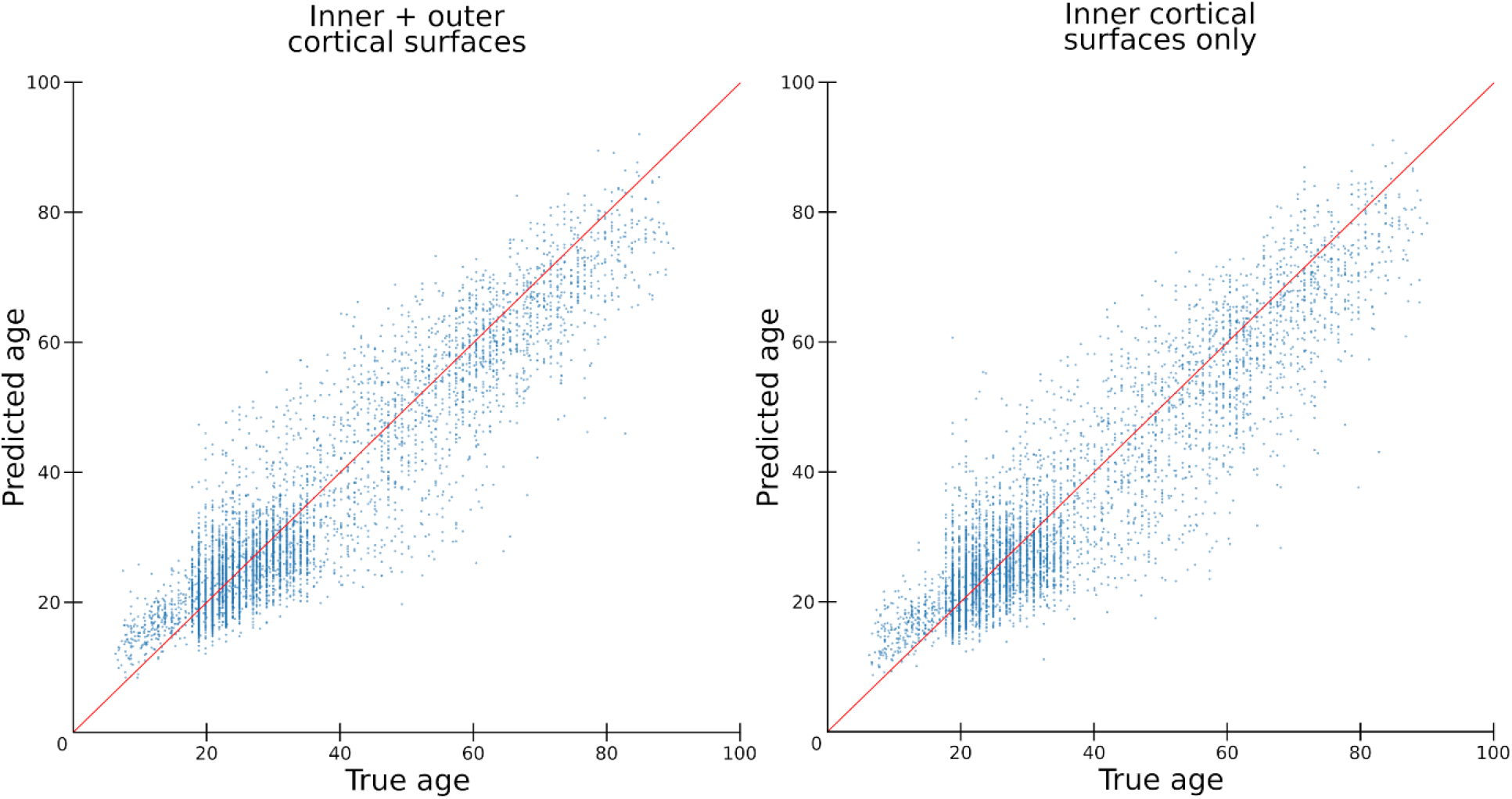
Results of age prediction with all the cortical surfaces or only the inner cortical surfaces. When all cortical surfaces are used, the Pearson’s coefficient of correlation *(R)* between the predicted and actual ages is *R* = 0.93. When only the inner cortical surfaces are used, *R*=0.92.

#### 3.1.2.2 Mapping discriminative brain regions

When using both the bilateral inner cortical surfaces only (i.e., only using morphologic features of the brain’s sulcal/gyral folding pattern), or the bilateral inner and outer cortical surfaces (i.e., morphologic features of the sulcal/gyral folding pattern and information regarding cortical thickness), there is a general trend towards the temporal and parietal lobes in younger subjects with a progressive inclusion of the frontal lobes in older subjects. Supplemental video 1 shows the most influential brain regions for age prediction over Gaussian sliding window (standard deviation 4 years) from 10 to 90 years old. Figure 9 shows three snapshots of this video at ages 20, 40 and 60 years old.

**Figure 9.**
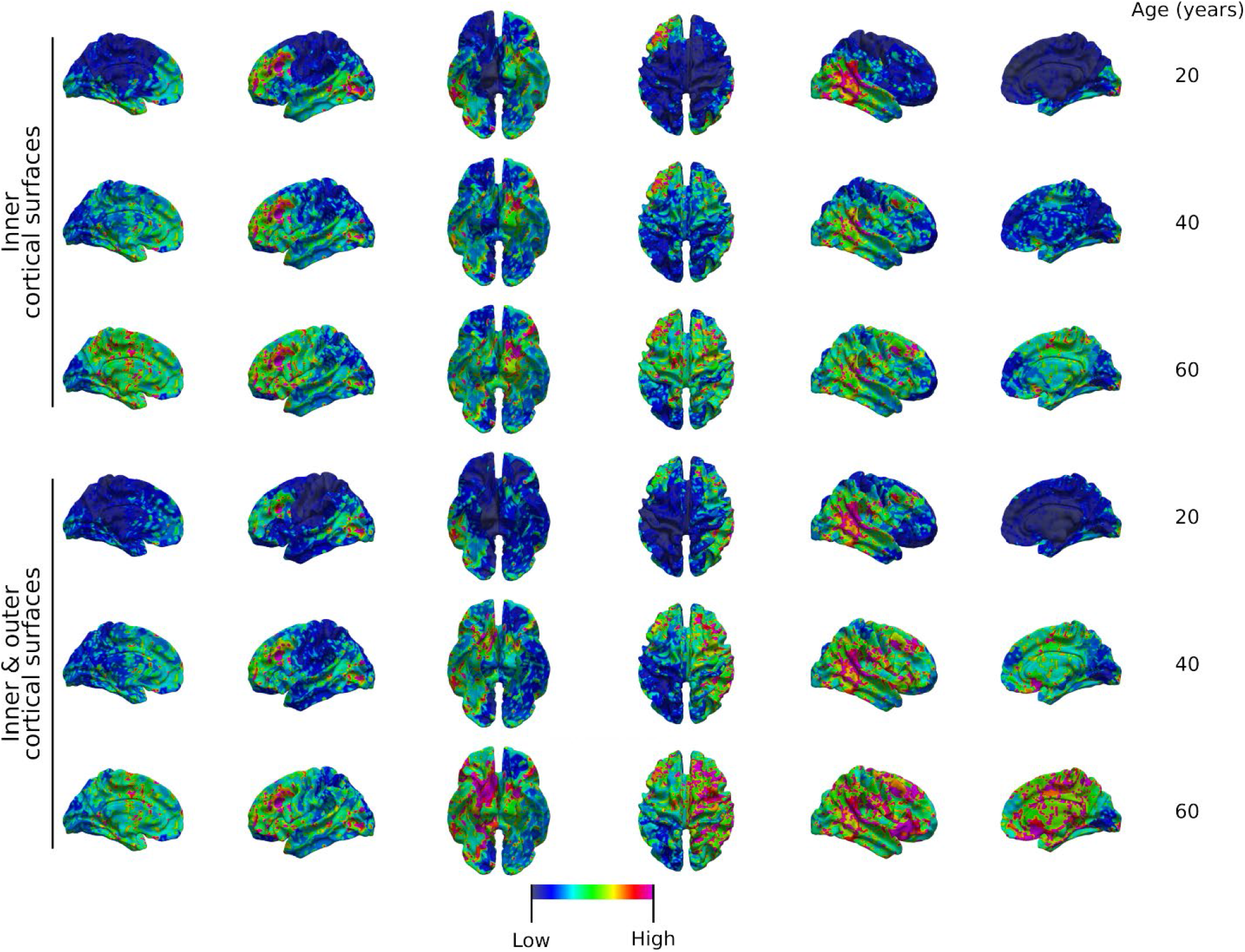
Regression activations maps (RAM) at ages 20, 40 and 60 years using the inner cortical surfaces, or both the inner and outer cortical surfaces. The color indicates cortical areas with either low or high involvement in the decision.

#### 3.1.2.3 Brain regions involved in correct prediction of youth (10-25 years)

In addition to this overall trend, younger subjects tended to be predicted correctly based on brain morphology alone (bilateral inner cortical surfaces only) primarily at the following brain regions: left medial prefrontal cortex, left dorsolateral prefrontal cortex, left inferior parietal lobule, left superior and inferior temporal gyri, right inferior parietal lobule (strong) and right occipital pole. Using a combination of brain morphology information and cortical thickness (bilateral inner and outer cortical surfaces), the left dorsolateral prefrontal cortex, left inferior parietal lobule, right inferior parietal lobule (strong) and right middle frontal gyri tended to be the most accurately predictive of young age.

#### 3.1.2.4 Brain regions involved in the correct prediction of adulthood (30-60 years)

Adult subjects, tended to be predicted correctly based on brain morphology alone primarily at the following brain regions: left medial paracentral lobule, left dorsolateral prefrontal cortex (to a lesser degree than in youth), left orbitofrontal cortex and the left prefrontal cortex including superior, middle and inferior frontal gyri). Using a combination of brain morphology and cortical thickness, the left medial paracentral lobule, left inferior parietal lobule, right greater than left orbitofrontal cortex and entire right frontal, temporal and parietal corticies over the cerebral convexity were found to be the most accurately predictive of adult age.

## 4.1 Discussion

### 4.1.1 An efficient surface-based method for cortical shape analysis

This study introduces a novel deep learning approach for the analysis of cortical morphology. It relies on cortical surface models, obtained from T1-weighted MR images, which are best suited to represent the folded sheet nature of the cortical ribbon (Dale, Fischl et al. 1999, Fischl, Sereno et al. 1999). This constitutes a significant departure from the classic volumetric analysis of brain imaging data (Bernal, Kushibar et al. 2019). Overall, the method uses graph convolutional neural networks to learn from data organized on non-Euclidean support, as it is the case for surface models, and benefits of the same advantages as traditional CNNs. In particular, this alleviates the challenging need to design relevant cortical geometric features tailored to a specific application. Using a large set of 6410 healthy subjects, we showed that this approach successfully predicted subjects’ age and sex form cortical folding patterns in a data-driven way. Importantly, our network performed well even using only the inner cortical surfaces, which models the grey/white interface therefore blinding the network to potential useful information such as cortical thickness. Using this approach, the accuracy of sex prediction was 85.23% and predicted age was in excellent agreement with the actual age with a Pearson’s coefficient of correlation (R) equal to 0.92. We also trained networks using both the inner and outer cortical surfaces, adding the possibility to the network to infer anatomic features of the inner surface relative to the outer surface, such as cortical thickness. In this situation, the performance accuracy was 87.99% for sex prediction and the correlation coefficient between predicted and actual ages was equal to 0.93, showing that adding the outer cortical surface only modestly increased the accuracy of the predictions and that cortical folding embeds a rich information.

These results are consistent with previous T1-weighted MRI based studies, where sex prediction accuracy ranged between 0.78 and 0.89 (Wachinger, Golland et al. 2015, Nieuwenhuis, Schnack et al. 2017, Pinaya, Mechelli et al. 2018), and age prediction reached R = 0.90 – 0.96 (Franke, Ziegler et al. 2010, Wachinger, Golland et al. 2015, Valizadeh, Hänggi et al. 2016, Gutierrez Becker, Klein et al. 2018). Our results extend these previous findings and are therefore notable given the increased complexity and heterogeneity of our dataset including our inclusion of children and 1888 subjects for whom age was imprecisely provided with a 2- or 5-year age bin rather than exact subject age, both of which add additional challenges to network training and resulting accuracy. In addition, we included data from 13 different cohorts of subjects from around the world using variable imaging systems and acquisition parameters. Therefore, the accuracy levels we report further emphasize the robustness and generalizability of this surface-based graph convolutional neural network approach.

Another advantage of surface-based deep learning is the natural two-dimensional representation of the cortical sheet, which is not only better suited to describe cortical topology but also helps to massively reduce the dimensionality of the data. In this study, the cortical ribbon was accurately modeled with 20,484 points, which represents fewer data points than a cubic grid of size 28^3^ and almost a 400-fold dimensionality reduction compared to standard whole brain 3D image of size 200^3^. Dimensionality reduction is important in machine learning applications as it helps mitigate the “curse of dimension”, minimizes undesired statistical properties associated with high dimensional data (Jimenez and Landgrebe 1998) and can significantly decrease the computational burden during the learning process.

### 4.1.2 Predictive mapping

The advantages of a CNN, and of a gCNN in particular, approach to brain imaging analysis includes its ability to produce topography distribution of brain areas the most involved classification and regression tasks. Despite deep learning’s many notable strengths, one of its most significant drawbacks has been the inability to interpret the methods used by the network to make an accurate prediction. gCNN analysis offers the ability to circumvent this to some degree in its ability to generate activation maps following correct prediction by the network. These maps can be generated at the subject or group level. When produced for a group, one can begin to understand the brain regions that were found to play a role in the correct prediction of the task at hand (in this case, correct prediction of age and sex), thus providing basic neuroscientific insights into the network’s decision making process. The characteristics themselves of the brain regions that contributed to successful prediction of age or sex remain unknown but one can begin to understand the distribution of brain regions involved in correct prediction. Said another way, these activation maps can provide information about which brain regions contributed to correct prediction but cannot provide information about what the features were in those locations that contributed to that correct prediction.

Class activation maps (CAMs) produced for correct sex prediction further inform the current understanding of sexual oligomorphism in the human brain. The frontal and parietal lobes were found to contain features predictive of female sex whereas male sex was more predicted by features within the bilateral perirolandic and antero-medial temporal structures (Figure 7). These findings recapitulate the work of many other groups who have defined a variety of cortical features including cortical thickness, cortical complexity, cortical gradient and fractal dimensionality, among others within these brain regions as variable across the human sexes (Zilles, Armstrong et al. 1988, Nopoulos, Flaum et al. 2000, Im, Lee et al. 2006, Luders, Thompson et al. 2006, Sowell, Peterson et al. 2007, Luders, Narr et al. 2008, Awate, Yushkevich et al. 2009, Salat, Lee et al. 2009, Awate, Yushkevich et al. 2010, Luders and Toga 2010, Lv, Li et al. 2010, Creze, Versheure et al. 2014).

Regression activation maps (RAMs) produced for correct age prediction demonstrated similarly consistent findings with previous literature. Specifically, the occipital lobes, frontal poles and entorhinal cortices were found to be more strongly predictive of youth with an ever expanding involvement of the frontal and parietal lobes in the accurate prediction of advancing age (Figure 9). This predominant role of the frontal, fronto-basal and occipital cortices in youth has been described before (Gogtay, Giedd et al. 2004, Kochunov, Mangin et al. 2005, Lemaître, Crivello et al. 2005, Shaw, Greenstein et al. 2006, Sowell, Peterson et al. 2007, Toro, Perron et al. 2008, Salat, Lee et al. 2009, Liu, Wen et al. 2010, Westlye, Walhovd et al. 2010, McGinnis, Brickhouse et al. 2011, Lemaitre, Goldman et al. 2012, Madan and Kensinger 2016, Jockwitz, Caspers et al. 2017). Similarly, our findings of a posterior toward anterior gradient of cortical influence in the aging process has also previously been described (Salat, Lee et al. 2009, Westlye, Walhovd et al. 2010). Together, these findings support the notion that gCNN analysis is able to identify morphologic features using only brain surface data to accurately predict demographic characteristics.

### 4.1.3 Conclusions

Here we demonstrate the accurate prediction of demographic features using a geometric deep learning approach on the brain’s shape. Beyond the currently predicted age and sex, the described method offers a wide range of potential applications including exploring the role of brain age in the aging process, understanding how brain age correlates with functional performance across the developmental spectrum, understanding sex-related brain features in the context of the gender identity spectrum as well as computer-aided detection of subtle brain pathologies that may escape visual identification, among others. The relevance of this report is not only in the accuracy rates achieved which are roughly equivalent to other published methods but rather in the ability of a novel computational tool to extract meaningful characteristics from the brain’s sulcal/gyral morphology alone.

## 6.1 Acknowledgements

Data were provided in part by the Brain Genomics Superstruct Project of Harvard University and the Massachusetts General Hospital, (Principal Investigators: Randy Buckner, Joshua Roffman, and Jordan Smoller), with support from the Center for Brain Science Neuroinformatics Research Group, the Athinoula A. Martinos Center for Biomedical Imaging, and the Center for Human Genetic Research. 20 individual investigators at Harvard and MGH generously contributed data to the overall project. Data were provided in part by the Human Connectome Project, WU-Minn Consortium (Principal Investigators: David Van Essen and Kamil Ugurbil; 1U54MH091657) funded by the 16 NIH Institutes and Centers that support the NIH Blueprint for Neuroscience Research; and by the McDonnell Center for Systems Neuroscience at Washington University. Part of this data was obtained from the OpenfMRI database. Its accession number is ds000221.

This research was supported in part through the computational resources and staff contributions provided for the Quest high performance computing facility at Northwestern University which is jointly supported by the Office of the Provost, the Office for Research, and Northwestern University Information Technology.

## Funding Sources

This research did not receive any specific grant from funding agencies in the public, commercial or not-for-profit sectors.

## Notes

### Competing Interest Statement

The authors have declared no competing interest.

## References

Awate, S. P., P. Yushkevich, D. Licht and J. C. Gee (2009). “Gender differences in cerebral cortical folding: multivariate complexity-shape analysis with insights into handling brain-volume differences.” Med Image Comput Comput Assist Interv 12(Pt 2): 200–207.

Awate, S. P., P. A. Yushkevich, Z. Song, D. J. Licht and J. C. Gee (2010). “Cerebral cortical folding analysis with multivariate modeling and testing: Studies on gender differences and neonatal development.” Neuroimage 53(2): 450–459.

Bernal, J., K. Kushibar, D. S. Asfaw, S. Valverde, A. Oliver, R. Martí and X. Lladó (2019). “Deep convolutional neural networks for brain image analysis on magnetic resonance imaging: a review.” Artificial Intelligence in Medicine 95: 64–81.

Besson, P., F. Andermann, F. Dubeau and A. Bernasconi (2008). “Small focal cortical dysplasia lesions are located at the bottom of a deep sulcus.” Brain 131(12): 3246–3255.

Bronstein MM B. J., LeCun Y, Szlam A, & Vandergheynst P (2017). “Geometric Deep Learning: Going beyond Euclidean data.” Ieee Signal Proc Mag 34(4): 18–42.

Cachia, A., M.-L. Paillère-Martinot, A. Galinowski, D. Januel, R. de Beaurepaire, F. Bellivier, E. Artiges, J. Andoh, D. Bartrés-Faz, E. Duchesnay, D. Rivière, M. Plaze, J.-F. Mangin and J.-L. Martinot (2008). “Cortical folding abnormalities in schizophrenia patients with resistant auditory hallucinations.” NeuroImage 39(3): 927–935.

Cachia, A., M. Roell, J.-F. Mangin, Z. Y. Sun, A. Jobert, L. Braga, O. Houde, S. Dehaene and G. Borst (2018). “How interindividual differences in brain anatomy shape reading accuracy.” Brain Structure and Function 223(2): 701–712.

Caviness, V. S. (1975). “Mechanical model of brain convolutional development.” Science 189(4196): 18.

Creze, M., L. Versheure, P. Besson, C. Sauvage, X. Leclerc and P. Jissendi-Tchofo (2014). “Age- and gender-related regional variations of human brain cortical thickness, complexity, and gradient in the third decade.” Hum Brain Mapp 35(6): 2817–2835.

Dale, A. M., B. Fischl and M. I. Sereno (1999). “Cortical Surface-Based Analysis: I. Segmentation and Surface Reconstruction.” NeuroImage 9(2): 179–194.

Defferrard, M., X. Bresson and P. Vandergheynst (2016). Convolutional neural networks on graphs with fast localized spectral filtering. Advances in Neural Information Processing Systems.

Dubois, J., M. Benders, C. Borradori-Tolsa, A. Cachia, F. Lazeyras, R. Ha-Vinh Leuchter, S. V. Sizonenko, S. K. Warfield, J. F. Mangin and P. S. Hüppi (2008). “Primary cortical folding in the human newborn: an early marker of later functional development.” Brain 131(8): 2028–2041.

Fischl, B., N. Rajendran, E. Busa, J. Augustinack, O. Hinds, B. T. T. Yeo, H. Mohlberg, K. Amunts and K. Zilles (2008). “Cortical Folding Patterns and Predicting Cytoarchitecture.” Cerebral Cortex 18(8): 1973–1980.

Fischl, B., M. I. Sereno and A. M. Dale (1999). “Cortical Surface-Based Analysis: II: Inflation, Flattening, and a Surface-Based Coordinate System.” NeuroImage 9(2): 195–207.

Fischl, B., M. I. Sereno, R. B. Tootell and A. M. Dale (1999). “High-resolution intersubject averaging and a coordinate system for the cortical surface.” Hum Brain Mapp 8(4): 272–284.

Franke, K., G. Ziegler, S. Klöppel and C. Gaser (2010). “Estimating the age of healthy subjects from T1-weighted MRI scans using kernel methods: Exploring the influence of various parameters.” NeuroImage 50(3): 883–892.

Gogtay, N., J. N. Giedd, L. Lusk, K. M. Hayashi, D. Greenstein, A. C. Vaituzis, T. F. Nugent, D. H. Herman, L. S. Clasen, A. W. Toga, J. L. Rapoport and P. M. Thompson (2004). “Dynamic mapping of human cortical development during childhood through early adulthood.” Proc Natl Acad Sci U S A 101(21): 8174–8179.

Gutierrez Becker, B., T. Klein and C. Wachinger (2018). “Gaussian process uncertainty in age estimation as a measure of brain abnormality.” NeuroImage 175: 246–258.

Hammond, D. K., P. Vandergheynst and R. Gribonval (2011). “Wavelets on graphs via spectral graph theory.” Applied and Computational Harmonic Analysis 30(2): 129–150.

He, K. M., X. Y. Zhang, S. Q. Ren and J. Sun (2016). “Deep Residual Learning for Image Recognition.” 2016 Ieee Conference on Computer Vision and Pattern Recognition (Cvpr): 770–778.

Hilgetag, C. and H. Barbas (2005). “Developmental mechanics of the primate cerebral cortex.” Anatomy and Embryology 210(5-6): 411–417.

Im, K., J.-M. Lee, U. Yoon, Y.-W. Shin, S. B. Hong, I. Y. Kim, J. S. Kwon and S. I. Kim (2006). “Fractal dimension in human cortical surface: Multiple regression analysis with cortical thickness, sulcal depth, and folding area.” Human Brain Mapping 27(12): 994–1003.

Im, K., J. M. Lee, J. Lee, Y. W. Shin, I. Y. Kim, J. S. Kwon and S. I. Kim (2006). “Gender difference analysis of cortical thickness in healthy young adults with surface-based methods.” Neuroimage 31(1): 31–38.

Jimenez, L. O. and D. A. Landgrebe (1998). “Supervised classification in high-dimensional space: geometrical, statistical, and asymptotical properties of multivariate data.” IEEE Transactions on Systems, Man, and Cybernetics, Part C (Applications and Reviews) 28(1): 39–54.

Jockwitz, C., S. Caspers, S. Lux, K. Jütten, A. Schleicher, S. B. Eickhoff, K. Amunts and K. Zilles (2017). “Age- and function-related regional changes in cortical folding of the default mode network in older adults.” Brain Struct Funct 222(1): 83–99.

Kingma, D. P. and J. J. a. p. a. Ba (2014). “Adam: A method for stochastic optimization.”

Kochunov, P., J.-F. Mangin, T. Coyle, J. Lancaster, P. Thompson, D. Rivière, Y. Cointepas, J. Régis, A. Schlosser, D. R. Royall, K. Zilles, J. Mazziotta, A. Toga and P. T. Fox (2005). “Age-related morphology trends of cortical sulci.” Human Brain Mapping 26(3): 210–220.

Kochunov, P., J. F. Mangin, T. Coyle, J. Lancaster, P. Thompson, D. Rivière, Y. Cointepas, J. Régis, A. Schlosser, D. R. Royall, K. Zilles, J. Mazziotta, A. Toga and P. T. Fox (2005). “Age-related morphology trends of cortical sulci.” Hum Brain Mapp 26(3): 210–220.

Kroenke, C. D. and P. V. Bayly (2018). “How Forces Fold the Cerebral Cortex.” The Journal of Neuroscience 38(4): 767.

LeCun, Y., Y. Bengio and G. Hinton (2015). “Deep learning.” Nature 521: 436.

Lemaitre, H., A. L. Goldman, F. Sambataro, B. A. Verchinski, A. Meyer-Lindenberg, D. R. Weinberger and V. S. Mattay (2012). “Normal age-related brain morphometric changes: nonuniformity across cortical thickness, surface area and gray matter volume?” Neurobiol Aging 33(3): 617.e611–619.

Lemaître, H., F. Crivello, B. Grassiot, A. Alpérovitch, C. Tzourio and B. Mazoyer (2005). “Age- and sex-related effects on the neuroanatomy of healthy elderly.” Neuroimage 26(3): 900–911.

Liu, T., W. Wen, W. Zhu, J. Trollor, S. Reppermund, J. Crawford, J. S. Jin, S. Luo, H. Brodaty and P. Sachdev (2010). “The effects of age and sex on cortical sulci in the elderly.” Neuroimage 51(1): 19–27.

Luders, E., K. L. Narr, R. M. Bilder, P. R. Szeszko, M. N. Gurbani, L. Hamilton, A. W. Toga and C. Gaser (2008). “Mapping the relationship between cortical convolution and intelligence: effects of gender.” Cereb Cortex 18(9): 2019–2026.

Luders, E., P. M. Thompson, K. L. Narr, A. W. Toga, L. Jancke and C. Gaser (2006). “A curvature-based approach to estimate local gyrification on the cortical surface.” Neuroimage 29(4): 1224–1230.

Luders, E. and A. W. Toga (2010). “Sex differences in brain anatomy.” Prog Brain Res 186: 3–12.

Lv, B., J. Li, H. He, M. Li, M. Zhao, L. Ai, F. Yan, J. Xian and Z. Wang (2010). “Gender consistency and difference in healthy adults revealed by cortical thickness.” Neuroimage 53(2): 373–382.

Madan, C. R. and E. A. Kensinger (2016). “Cortical complexity as a measure of age-related brain atrophy.” Neuroimage 134: 617–629.

McGinnis, S. M., M. Brickhouse, B. Pascual and B. C. Dickerson (2011). “Age-related changes in the thickness of cortical zones in humans.” Brain Topogr 24(3-4): 279–291.

Nieuwenhuis, M., H. G. Schnack, N. E. van Haren, J. Lappin, C. Morgan, A. A. Reinders, D. Gutierrez-Tordesillas, R. Roiz-Santiañez, M. S. Schaufelberger, P. G. Rosa, M. V. Zanetti, G. F. Busatto, B. Crespo-Facorro, P. D. McGorry, D. Velakoulis, C. Pantelis, S. J. Wood, R. S. Kahn, J. Mourao-Miranda and P. Dazzan (2017). “Multi-center MRI prediction models: Predicting sex and illness course in first episode psychosis patients.” NeuroImage 145: 246-253.

Nopoulos, P., M. Flaum, D. O’Leary and N. C. Andreasen (2000). “Sexual dimorphism in the human brain: evaluation of tissue volume, tissue composition and surface anatomy using magnetic resonance imaging.” Psychiatry Res 98(1): 1–13.

Nordahl, C. W., D. Dierker, I. Mostafavi, C. M. Schumann, S. M. Rivera, D. G. Amaral and D. C. Van Essen (2007). “Cortical Folding Abnormalities in Autism Revealed by Surface-Based Morphometry.” The Journal of Neuroscience 27(43): 11725.

Penttilä, J., M.-L. Paillère-Martinot, J.-L. Martinot, D. Ringuenet, M. Wessa, J. Houenou, T. Gallarda, F. Bellivier, A. Galinowski, P. Bruguière, F. Pinabel, M. Leboyer, J.-P. Olié, E. Duchesnay, E. Artiges, J.-F. Mangin and A. Cachia (2009). “Cortical folding in patients with bipolar disorder or unipolar depression.” Journal of psychiatry & neuroscience: JPN 34(2): 127–135.

Pinaya, W. H. L., A. Mechelli and J. R. Sato (2018). “Using deep autoencoders to identify abnormal brain structural patterns in neuropsychiatric disorders: A large-scale multi-sample study.” Human Brain Mapping 0(0).

Rakic, P. (1988). “Specification of cerebral cortical areas.” Science 241(4862): 170.

Salat, D. H., S. Y. Lee, A. J. van der Kouwe, D. N. Greve, B. Fischl and H. D. Rosas (2009). “Age-associated alterations in cortical gray and white matter signal intensity and gray to white matter contrast.” Neuroimage 48(1): 21–28.

Schaer, M., M. B. Cuadra, L. Tamarit, F. Lazeyras, S. Eliez and J. Thiran (2008). “A Surface-Based Approach to Quantify Local Cortical Gyrification.” IEEE Transactions on Medical Imaging 27(2): 161–170.

Schmitgen, M. M., M. S. Depping, C. Bach, N. D. Wolf, K. M. Kubera, N. Vasic, D. Hirjak, F. Sambataro and R. C. Wolf (2019). “Aberrant cortical neurodevelopment in major depressive disorder.” Journal of Affective Disorders 243: 340–347.

Shaw, P., D. Greenstein, J. Lerch, L. Clasen, R. Lenroot, N. Gogtay, A. Evans, J. Rapoport and J. Giedd (2006). “Intellectual ability and cortical development in children and adolescents.” Nature 440(7084): 676–679.

Sherif, T., P. Rioux, M.-E. Rousseau, N. Kassis, N. Beck, R. Adalat, S. Das, T. Glatard and A. C. Evans (2014). “CBRAIN: a web-based, distributed computing platform for collaborative neuroimaging research.” 8(54).

Shuman, D. I., S. K. Narang, P. Frossard, A. Ortega and P. Vandergheynst (2013). “The Emerging Field of Signal Processing on Graphs.” Ieee Signal Processing Magazine 30(3): 83–98.

Sowell, E. R., B. S. Peterson, E. Kan, R. P. Woods, J. Yoshii, R. Bansal, D. Xu, H. Zhu, P. M. Thompson and A. W. Toga (2007). “Sex differences in cortical thickness mapped in 176 healthy individuals between 7 and 87 years of age.” Cereb Cortex 17(7): 1550–1560.

Thompson, P. M., A. D. Lee, R. A. Dutton, J. A. Geaga, K. M. Hayashi, M. A. Eckert, U. Bellugi, A. M. Galaburda, J. R. Korenberg, D. L. Mills, A. W. Toga and A. L. Reiss (2005). “Abnormal Cortical Complexity and Thickness Profiles Mapped in Williams Syndrome.” The Journal of Neuroscience 25(16): 4146.

Toro, R., M. Perron, B. Pike, L. Richer, S. Veillette, Z. Pausova and T. Paus (2008). “Brain Size and Folding of the Human Cerebral Cortex.” Cerebral Cortex 18(10): 2352–2357.

Valizadeh, S. A., J. Hänggi, S. Mérillat and L. Jäncke (2016). “Age prediction on the basis of brain anatomical measures.” Human Brain Mapping 38(2): 997–1008.

Van Essen, D. C. (1997). “A tension-based theory of morphogenesis and compact wiring in the central nervous system.” Nature 385(6614): 313–318.

Wachinger, C., P. Golland, W. Kremen, B. Fischl and M. Reuter (2015). “BrainPrint: A discriminative characterization of brain morphology.” NeuroImage 109: 232–248.

Wang, Z. and J. J. a. p. a. Yang (2017). “Diabetic Retinopathy Detection via Deep Convolutional Networks for Discriminative Localization and Visual Explanation.”

Westlye, L. T., K. B. Walhovd, A. M. Dale, A. Bjørnerud, P. Due-Tønnessen, A. Engvig, H. Grydeland, C. K. Tamnes, Y. Østby and A. M. Fjell (2010). “Differentiating maturational and aging-related changes of the cerebral cortex by use of thickness and signal intensity.” Neuroimage 52(1): 172–185.

Whittle, S., N. B. Allen, A. Fornito, D. I. Lubman, J. G. Simmons, C. Pantelis and M. Yücel (2009). “Variations in cortical folding patterns are related to individual differences in temperament.” Psychiatry Research: Neuroimaging 172(1): 68–74.

Zhou, B., A. Khosla, A. Lapedriza, A. Oliva and A. Torralba (2016). “Learning Deep Features for Discriminative Localization.” 2016 Ieee Conference on Computer Vision and Pattern Recognition (Cvpr): 2921–2929.

Zilles, K. and K. Amunts (2010). “Centenary of Brodmann&s map — conception and fate.” Nature Reviews Neuroscience 11: 139.

Zilles, K., E. Armstrong, A. Schleicher and H. J. Kretschmann (1988). “The human pattern of gyrification in the cerebral cortex.” Anat Embryol (Berl) 179(2): 173–179.

