## Supplemental Information for "Geometric deep learning on brain shape predicts sex and age"

#### Datasets Included

- Autism Brain Imaging Data Exchange II (ABIDE II)
- Age-ility
- Cambridge Centre for Ageing Neuroscience (CamCan)
- Consortium for Reliability and Reproducibility (CoRR)
- Dallas Lifespan Brain Study (DLBS)
- Brain Genomics Superstruct Project (GSP)
- Human Connectome Project (HCP)
- Information eXtraction from Images (IXI)
- MPI-Leipzig Mind Brain Body (MPI-LMBB)
- Enhanced Nathan Kline Institute-Rockland Sample (NKI-RS)
- Southwest University Adult Lifespan Dataset (SALD)

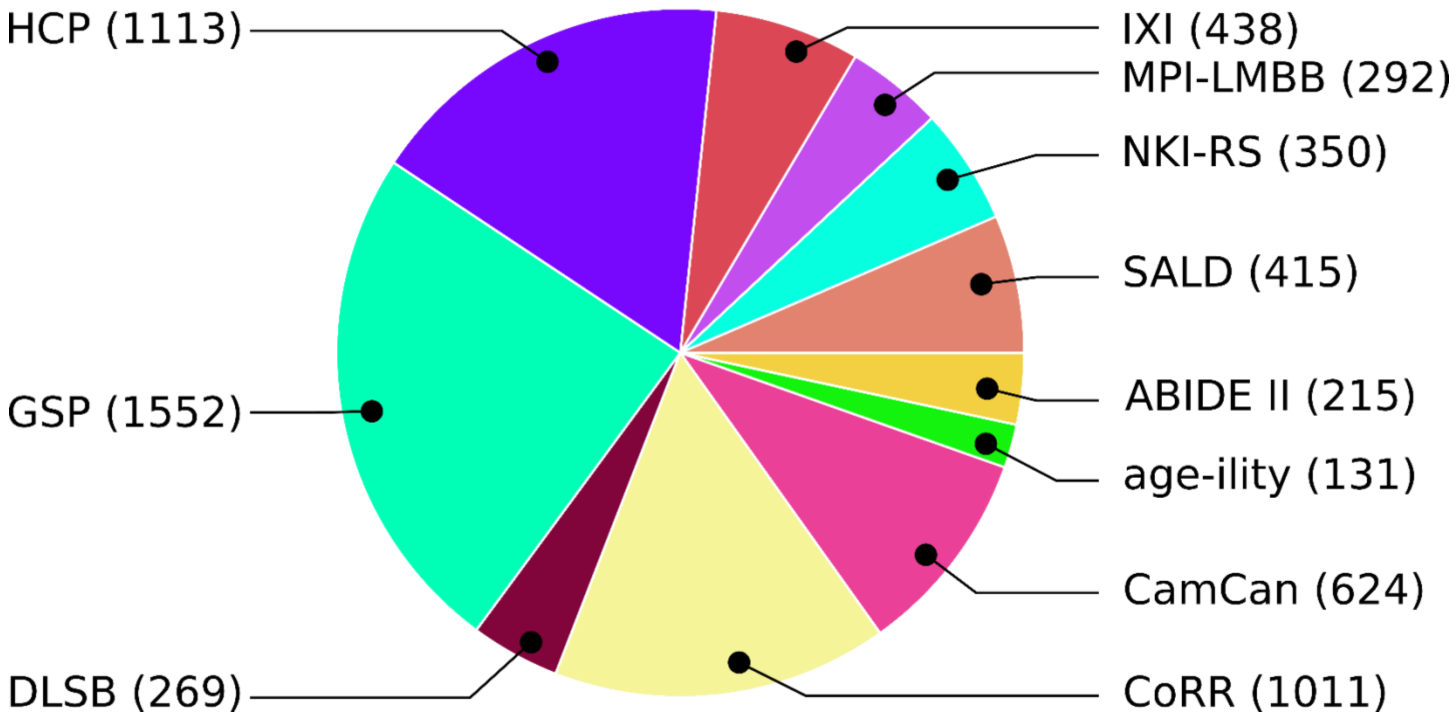

#### Autism Brain Imaging Data Exchange II (ABIDE II)

**Final number of healthy subjects included in the study: 215**

*Website:* http://fcon_1000.projects.nitrc.org/indi/abide/abide_II.html

*Description:* The establishment of ABIDE I demonstrated the feasibility and utility of aggregating MRI data across sites. However, the complexity of the connectome, along with the substantial heterogeneity of Autism Spectrum Disorder (ASD) and initial results from ABIDE I data analyses underscore the need for even larger and better-characterized samples. Accordingly, with support of an award from the National Institute of Mental Health (R21MH107045), ABIDE II was established to further promote discovery science on the brain connectome in ASD. To date, ABIDE II has aggregated over 1000 additional datasets with greater phenotypic characterization, particularly in regard to measures of core ASD and associated symptoms. In addition, two collections include longitudinal samples of data collected from 38 individuals at two time points (1-4 year interval). To date, ABIDE II involves 19 sites - ten charter institutions and seven new members - overall donating 1114 datasets from 521 individuals with ASD and 593 controls (age range: 5-64 years). These data have been openly released to the scientific community on June 2016. In accordance with HIPAA guidelines and 1000 Functional Connectomes Project / INDI protocols, all datasets are anonymous, with no protected health information included.

*Image acquisition:* ABIDE II groups data collected from different centers. Acquisition parameters and number of healthy subjects included are the following:

1. **Barrow Neurological Institute (BNI)**
   *Detailed acquisition parameters:*
   http://fcon_1000.projects.nitrc.org/indi/abide/scan_params/ABIDEII-BNI_1_scantable.pdf
   *Summary:* Images acquired on a 3-Tesla Philips Ingenia scanner, 15-channel headcoil. MPRAGE anatomic T1-weighted sequence (FA = 9°, TI = 900 ms, TE = shortest, in-plane resolution = 1.1 × 1.1 mm$^2$, slice thickness = 1.2 mm, slice gap = 0 mm, FOV = 270 × 252 mm$^2$, number of slices = 170)
   *Number of subjects included from this dataset:* 29
2. **Erasmus University Medical Center Rotterdam (EMC)**
   *Detailed acquisition parameters:*
   http://fcon_1000.projects.nitrc.org/indi/abide/scan_params/ABIDEII-EMC_1_scantable.pdf
   *Summary:* Images acquired on a 3-Tesla GE MR750 scanner, 8-channel headcoil. IR-FSPGR anatomic T1-weighted sequence (FA = 16°, TI = 350 ms, TE = 4.236 ms, in-plane resolution = 0.9 × 0.9 mm$^2$, slice thickness = 0.9 mm, slice gap = 0 mm, FOV = 230 × 230 mm$^2$, number of slices = 186)
   *Number of subjects included from this dataset:* 19
3. **ETH Zürich (ETH)**
   *Detailed acquisition parameters:*
   http://fcon_1000.projects.nitrc.org/indi/abide/scan_params/ABIDEII-ETH_1_scantable.pdf
   *Summary:* Images acquired on a 3-Tesla Philips Achieva scanner, 32-channel headcoil. Axial TFE anatomic T1-weighted sequence (FA = 8°, TI = 1150 ms, TE = shortest, in-plane resolution = 0.9 × 0.9 mm$^2$, slice thickness = 0.9 mm, slice gap = 0.09 mm, FOV = 230 × 230 mm$^2$, number of slices = 180)
   *Number of subjects included from this dataset:* 23
4. **Georgetown University (GU)**
   *Detailed acquisition parameters:*
   http://fcon_1000.projects.nitrc.org/indi/abide/scan_params/ABIDEII-GU_1_scantable.pdf
   *Summary*: Images acquired on a 3-Tesla Siemens TriTim scanner, 12-channel headcoil. MPRAGE anatomic T1-weighted sequence (FA = 7°, TI = 1100 ms, TE = 3.5 ms, in-plane resolution = 1 × 1 mm$^2$, slice thickness = 1 mm, slice gap = 0.5 mm, FOV = 256 × 256 mm$^2$, number of slices = 176)
   *Number of subjects included from this dataset:* 27
5. **Indiana University (IU)**
   *Detailed acquisition parameters:*
   http://fcon_1000.projects.nitrc.org/indi/abide/scan_params/ABIDEII-IU_1_scantable.pdf

*Summary*: Images acquired on a 3-Tesla Siemens TriTim scanner, 32-channel headcoil. 3D TFL anatomic T1-weighted sequence (FA = 8°, TI = 1000 ms, TE = 2.3 ms, in-plane resolution = 0.7 × 0.7 mm$^2$, slice thickness = 0.7 mm, slice gap = 0.35 mm, FOV = 224 × 224 mm$^2$, number of slices = 256)
*Number of subjects included from this dataset:* 9

6. **Institut Pasteur and Robert Debré Hospital (IP)**
   *Detailed acquisition parameters:*
   http://fcon_1000.projects.nitrc.org/indi/abide/scan_params/ABIDEII-IP_1_scantable.pdf
   *Summary*: Images acquired on a 1.5-Tesla Philips Achieva scanner, 8-channel headcoil. MPRAGE anatomic T1-weighted sequence (FA = 30°, TI = NA, TE = 5.6 ms, TR = 25 ms, in-plane resolution = 1 × 1 mm$^2$, slice thickness = 1 mm, slice gap = 0 mm, FOV = 240 × 240 mm$^2$, number of slices = 170)
   *Number of subjects included from this dataset:* 16
7. **NYU Langone Medical Center: Sample 1 (NYU1)**
   *Detailed acquisition parameters:*
   http://fcon_1000.projects.nitrc.org/indi/abide/scan_params/ABIDEII-NYU_1_scantable.pdf
   *Summary*: Images acquired on a 3-Tesla Siemens Allegra scanner, 8-channel headcoil. 3D TFL anatomic T1-weighted sequence (FA = 7°, TI = 1100 ms, TE = 3.25 ms, in-plane resolution = 1.3 × 1.0 mm$^2$, slice thickness = 1.33 mm, slice gap = 0.665 mm, FOV = 256 × 256 mm$^2$, number of slices = 128)
   *Number of subjects included from this dataset:* 19
8. **Oregon Health and Science University (OHSU)**
   *Detailed acquisition parameters:*
   http://fcon_1000.projects.nitrc.org/indi/abide/scan_params/ABIDEII-OHSU_1_scantable.pdf
   *Summary*: Images acquired on a 3-Tesla Siemens TriTim scanner, 12-channel headcoil. 3D TFL anatomic T1-weighted sequence (FA = 10°, TI = 900 ms, TE = 3.58 ms, in-plane resolution = 1 × 1 mm$^2$, slice thickness = 1.1 mm, slice gap = 0.55 mm, FOV = 256 × 240 mm$^2$, number of slices = 160)
   *Number of subjects included from this dataset:* 16
9. **Trinity Centre for Health Sciences (TCD)**
   *Detailed acquisition parameters:*
   http://fcon_1000.projects.nitrc.org/indi/abide/scan_params/ABIDEII-TCD_1_scantable.pdf
   *Summary*: Images acquired on a 3-Tesla Philips Achieva scanner, 8-channel headcoil. MPRAGE anatomic T1-weighted sequence (FA = 8°, TI = 1150 ms, TE = 3.9 ms, in-plane resolution = 0.9 × 0.9 mm$^2$, slice thickness = 0.9 mm, slice gap = 0.09 mm, FOV = 230 × 230 mm$^2$, number of slices = 190)
   *Number of subjects included from this dataset:* 19
10. **San Diego State University (SDSU)**
    *Detailed acquisition parameters:*
    http://fcon_1000.projects.nitrc.org/indi/abide/scan_params/ABIDEII-SDSU_1_scantable.pdf
    *Summary*: Images acquired on a 3-Tesla GE MR750 scanner, 8-channel headcoil. 3D SPGR anatomic T1-weighted sequence (FA = 8°, TI = 600 ms, TE = 3.172 ms, in-plane resolution = 1 × 1 mm$^2$, slice thickness = 1 mm, slice gap = 1 mm, FOV = 256 × 256 mm$^2$, number of slices = 172)
    *Number of subjects included from this dataset:* 7
11. **Stanford University (SU)**
    *Detailed acquisition parameters:*
    http://fcon_1000.projects.nitrc.org/indi/abide/scan_params/ABIDEII-SU_2/anat.pdf
    *Summary*: Images acquired on a 3-Tesla GE Signa scanner. T1-weighted sequence (FA = 11°, TI = NA, TE = 1.8 ms, TR = 5.9 ms, in-plane resolution = 0.9375 × 0.9375 mm$^2$, slice thickness = 1 mm, slice gap = 0.25 mm, FOV = 240 × 240 mm$^2$, number of slices = 132)
    *Number of subjects included from this dataset:* 2
12. **University of California Davis (UCD)**
    *Detailed acquisition parameters:*
    http://fcon_1000.projects.nitrc.org/indi/abide/scan_params/ABIDEII-UCD_1_scantable.pdf

*Summary*: Images acquired on a 3-Tesla Siemens TriTim scanner, 32-channel headcoil. MPRAGE T1-weighted sequence (FA = 8°, TI = 1050 ms, TE = 3.16 ms, TR = 2000 ms, in-plane resolution = 1 × 1 $mm^2$, slice thickness = 1 mm, slice gap = 0.5 mm, FOV = 256 × 224 $mm^2$, number of slices = 192)
*Number of subjects included from this dataset:* 10

**13. University of California Los Angeles (UCLA)**
*Detailed acquisition parameters:*
http://fcon_1000.projects.nitrc.org/indi/abide/scan_params/ABIDEII-UCLA_1_scantable.pdf
*Summary*: Images acquired on a 3-Tesla Siemens TriTim scanner, 12-channel headcoil. MPRAGE T1-weighted sequence (FA = 9°, TI = 853 ms, TE = 2.86 ms, TR = 2300 ms, in-plane resolution = 1 × 1 $mm^2$, slice thickness = 1.2 mm, slice gap = 0.6 mm, FOV = 256 × 240 $mm^2$, number of slices = 160)
*Number of subjects included from this dataset:* 7

**14. University of Miami (MIA)**
*Detailed acquisition parameters:*
http://fcon_1000.projects.nitrc.org/indi/abide/scan_params/ABIDEII-U_MIA_1/anat.pdf
*Summary*: Images acquired on a 3-Tesla GE scanner. T1-weighted sequence (FA = 12°, TI = 650 ms, TE = NA, TR = NA, in-plane resolution = 1 × 1 $mm^2$, slice thickness = 1 mm, slice gap = NA, FOV = 216 × 256 $mm^2$, number of slices = 256)
*Number of subjects included from this dataset:* 5

**15. University of Utah School of Medicine (USM)**
*Detailed acquisition parameters:*
http://fcon_1000.projects.nitrc.org/indi/abide/scan_params/ABIDEII-USM_1_scantable.pdf
*Summary*: Images acquired on a 3-Tesla Siemens TriTim scanner, 12-channel headcoil. MPRAGE T1-weighted sequence (FA = 9°, TI = 900 ms, TE = 2.91 ms, TR = 900 ms, in-plane resolution = 1 × 1 $mm^2$, slice thickness = 1.2 mm, slice gap = 0.6 mm, FOV = 256 × 240 $mm^2$, number of slices = 160)
*Number of subjects included from this dataset:* 7

*Summary of final inclusion from ABIDE II:*

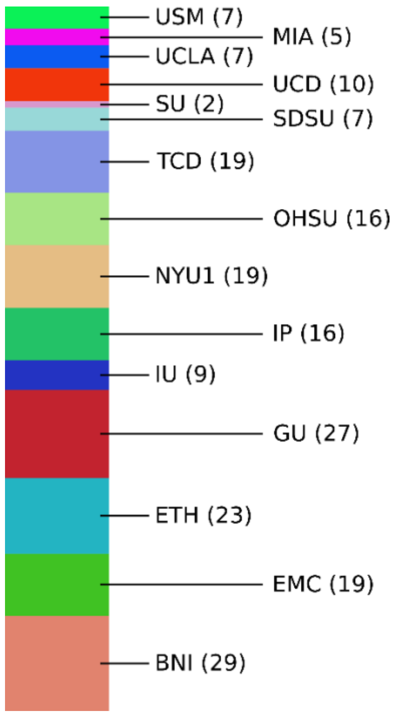

#### Age-ility

**Final number of healthy subjects included in the study: 131**

*Website:* https://www.nitrc.org/projects/age-ility/

*Description:* More information in the following publication (1).

*Image acquisition:* Images acquired on a 3-Tesla Siemens Skyra scanner, 32-channel headcoil. MPRAGE T1-weighted sequence (FA = 7°, TI = 1100 ms, TE = 3.5 ms, TR = 2200 ms, in-plane resolution = 1 × 1 mm2, slice thickness = 1 mm, slice gap = 0 mm, FOV = 256 × 256 $mm^2$, number of slices = 176)

#### Cambridge Centre for Ageing Neuroscience (CamCan)

**Final number of healthy subjects included in the study: 624**

*Website:* https://camcan-archive.mrc-cbu.cam.ac.uk/dataaccess/

*Description:* The Cambridge Centre for Ageing and Neuroscience (Cam-CAN) is a large-scale collaborative research project, launched in October 2010, with substantial funding from the Biotechnology and Biological Sciences Research Council (BBSRC). The Cam-CAN project is using epidemiological, behavioural, and neuroimaging data to understand how individuals can best retain cognitive abilities into old age.

This inventory contains details of data from Stage II of the Cambridge Centre for Ageing and Neuroscience (CamCAN) project. Here 700 adults were scanned using structural Magnetic Resonance Imaging (MRI), functional MRI (both resting and task-based), and magnetoencephalography (MEG) (resting and task-based). Many behavioural experiments were also performed outside of the scanner. The following page details all those datasets in the CC700 stage that are ready, as well as those that are still in the preprocessing pipeline. In time, data from Stages I and III will be released with more behavioural functional MRI and MEG scans.

The age-range of available participants is 18-88 years for each of the datasets below, approximately evenly distributed across seven decades. A detailed description of the cc700 dataset and pre-processing pipeline can be found in (2). A more general overview of the CamCAN dataset can be found in (3).

*Image acquisition:* Details of sequence parameters can be found:

https://camcan-archive.mrc-cbu.cam.ac.uk/dataaccess/pdfs/CAMCAN700_MR_params.pdf

Images acquired on a 3-Tesla Siemens TrioTim scanner, 32-channel headcoil. MPRAGE T1-weighted sequence (FA = 9°, TI = 900 ms, TE = 2.99 ms, TR = 2250 ms, in-plane resolution = 1 × 1 $mm^2$, slice thickness = 1 mm, slice gap = 0 mm, FOV = 256 × 240 $mm^2$, number of slices = 192)

#### Consortium for Reliability and Reproducibility (CoRR)

**Final number of healthy subjects included in the study: 1011**

*Website:* http://fcon_1000.projects.nitrc.org/indi/CoRR/html/

*Description:* The goal of CoRR was to create an open science resource for the imaging community that facilitates the assessment of test-retest reliability and reproducibility for functional and structural connectomics. In order to accomplish this, we have aggregated resting state fMRI (R-fMRI) and diffusion imaging data from laboratories around the world, and are sharing the data via the International Neuroimaging Data-sharing Initiative (INDI). This enables the:

- Establishment of test-retest reliability and reproducibility for commonly used MR-based connectome metrics
- Determination of the range of variation in the reliability and reproducibility of these metrics across imaging sites and retest study designs
- Creation of a standard/benchmark test-retest dataset for the evaluation of novel metrics

Given that this was a retrospective data collection, we have focused on basic phenotypic measures that are relatively standard in the neuroimaging field, as well as fundamental for analyses and sample characterization. Our phenotypic key is organized to reflect three classifications of variables: 1) core (i.e., minimal variables required to characterize any dataset), 2) preferred (i.e., variables that were strongly suggested for inclusion due to their relative import and/or likelihood of being collected by most sites), and 3) optional (variables that are data-set specific or only shared by a few sites).

*Image acquisition:* Overall, CoRR includes 33 datasets, of which 32 are currently available for download, consisting of 1629 subjects. Further details can be found in (4). Were included in this study:

1. **Berlin Mind and Brain (BMB1)**
   *Detailed acquisition parameters:*
   http://fcon_1000.projects.nitrc.org/indi/CoRR/html/_static/scan_parameters/BMB_1_scantable.pdf
   *Summary:* Images acquired on a 3-Tesla Siemens TrioTim scanner, 12-channel headcoil. 3D MPRAGE anatomic T1-weighted sequence (FA = 9°, TI = 900 ms, TE = 2.98 ms, TR = 2300 ms, in-plane resolution = 1 × 1 $mm^2$, slice thickness = 1 mm, slice gap = 0.5 mm, FOV = 256 × 256 $mm^2$, number of slices = 176)
   *Number of subjects included from this dataset:* 50
2. **Beijing Normal University (BNU1)**
   *Detailed acquisition parameters:*
   http://fcon_1000.projects.nitrc.org/indi/CoRR/html/_static/scan_parameters/BNU_1_scantable.pdf
   *Summary:* Images acquired on a 3-Tesla Siemens TrioTim scanner, 12-channel headcoil. 3D MPRAGE anatomic T1-weighted sequence (FA = 7°, TI = 1100 ms, TE = 3.39 ms, TR = 2530 ms, in-plane resolution = 1.3 × 1.0 $mm^2$, slice thickness = 1.3 mm, slice gap = 0.65 mm, FOV = 256 × 256 $mm^2$, number of slices = 144)
   *Number of subjects included from this dataset:* 32
3. **Beijing Normal University (BNU3)**
   *Detailed acquisition parameters:*
   http://fcon_1000.projects.nitrc.org/indi/CoRR/html/_static/scan_parameters/BNU_3_scantable.pdf
   *Summary:* Images acquired on a 3-Tesla Siemens TrioTim scanner, 12-channel headcoil. 3D MPRAGE anatomic T1-weighted sequence (FA = 7°, TI = 1100 ms, TE = 3.39 ms, TR = 2530 ms, in-plane resolution = 1.3 × 1.0 $mm^2$, slice thickness = 1.33 mm, slice gap = 0.6515 mm, FOV = 256 × 256 $mm^2$, number of slices = 128)
   *Number of subjects included from this dataset:* 34
4. **Hangzhou Normal University (HNU)**
   *Detailed acquisition parameters:*
   http://fcon_1000.projects.nitrc.org/indi/CoRR/html/_static/scan_parameters/HNU_1_scantable.pdf

*Summary:* Images acquired on a 3-Tesla GE Discovery MR750 scanner, 8-channel headcoil. 3D SPGR anatomic T1-weighted sequence (FA = 8°, TI = 450 ms, TE = min full, TR = 8.06 ms, in-plane resolution = 1.0 × 1.0 $mm^2$, slice thickness = 1.0 mm, slice gap = 0 mm, FOV = 250 × 250 $mm^2$, number of slices = 180)
*Number of subjects included from this dataset:* 29

5. **Institute of Automation, Chinese Academy of Sciences (IACAS)**
*Detailed acquisition parameters:*
http://fcon_1000.projects.nitrc.org/indi/CoRR/html/_static/scan_parameters/IACAS_1_scantable.pdf
*Summary:* Images acquired on a 3-Tesla GE Signa HDx scanner, 8-channel headcoil. 3D BRAVO anatomic T1-weighted sequence (FA = 7°, TI = 1100 ms, TE = 2.984 ms, TR = 7.788 ms, in-plane resolution = 1.0 × 1.0 $mm^2$, slice thickness = 1.0 mm, slice gap = 0 mm, FOV = 256 × 256 $mm^2$, number of slices = 192)
*Number of subjects included from this dataset:* 22

6. **Intrinsic Brain Activity, Test-Retest Dataset (IBATRT)**
*Detailed acquisition parameters:*
http://fcon_1000.projects.nitrc.org/indi/CoRR/html/_static/scan_parameters/IBATRT_scantable.pdf
*Summary:* Images acquired on a 3-Tesla Siemens TrioTim scanner, 12-channel headcoil. 3D MPRAGE anatomic T1-weighted sequence (FA = 8°, TI = 900 ms, TE = 3.02 ms, TR = 2600 ms, in-plane resolution = 1.0 × 1.0 $mm^2$, slice thickness = 1.0 mm, slice gap = 0.5 mm, FOV = 256 × 256 $mm^2$, number of slices = 176)
*Number of subjects included from this dataset:* 34

7. **Institute of Psychology, Chinese Academy of Sciences (IPCAS1)**
*Detailed acquisition parameters:*
http://fcon_1000.projects.nitrc.org/indi/CoRR/html/_static/scan_parameters/IPCAS_1_scantable.pdf
*Summary:* Images acquired on a 3-Tesla Siemens TrioTim scanner, 8-channel headcoil. 3D MPRAGE anatomic T1-weighted sequence (FA = 7°, TI = 1100 ms, TE = 2.51 ms, TR = 2530 ms, in-plane resolution = 1.0 × 1.0 $mm^2$, slice thickness = 1.3 mm, slice gap = 0.65 mm, FOV = 256 × 256 $mm^2$, number of slices = 128)
*Number of subjects included from this dataset:* 30

8. **Institute of Psychology, Chinese Academy of Sciences (IPCAS2)**
*Detailed acquisition parameters:*
http://fcon_1000.projects.nitrc.org/indi/CoRR/html/_static/scan_parameters/IPCAS_2_scantable.pdf
*Summary:* Images acquired on a 3-Tesla Siemens TrioTim scanner, 32-channel headcoil. 3D MPRAGE anatomic T1-weighted sequence (FA = 9°, TI = 900 ms, TE = 2.95 ms, TR = 2300 ms, in-plane resolution = 0.9 × 0.9 $mm^2$, slice thickness = 1.2 mm, slice gap = 0.6 mm, FOV = 240 × 240 $mm^2$, number of slices = 160)
*Number of subjects included from this dataset:* 29

9. **Institute of Psychology, Chinese Academy of Sciences (IPCAS3)**
*Detailed acquisition parameters:*
http://fcon_1000.projects.nitrc.org/indi/CoRR/html/_static/scan_parameters/IPCAS_3_scantable.pdf
*Summary:* Images acquired on a 3-Tesla Siemens TrioTim scanner, 8-channel headcoil. 3D MPRAGE anatomic T1-weighted sequence (FA = 7°, TI = 1100 ms, TE = 2.51 ms, TR = 2530 ms, in-plane resolution = 1.0 × 1.0 $mm^2$, slice thickness = 1.33 mm, slice gap = NA, FOV = 256 × 256 $mm^2$, number of slices = 128)
*Number of subjects included from this dataset:* 30

10. **Institute of Psychology, Chinese Academy of Sciences (IPCAS4)**
*Detailed acquisition parameters:*
http://fcon_1000.projects.nitrc.org/indi/CoRR/html/_static/scan_parameters/IPCAS_4_scantable.pdf
*Summary:* Images acquired on a 3-Tesla GE Discovery scanner, 8-channel headcoil. 3D SPGR anatomic T1-weighted sequence (FA = 8°, TI = 450 ms, TE = 3.136 ms, TR = 8068 ms, in-plane resolution = 1.0 × 1.0 $mm^2$, slice thickness = 1.0 mm, slice gap = 0 mm, FOV = 250 × 250 $mm^2$, number of slices = 250)
*Number of subjects included from this dataset:* 18

11. **Institute of Psychology, Chinese Academy of Sciences (IPCAS5)**
*Detailed acquisition parameters:*
http://fcon_1000.projects.nitrc.org/indi/CoRR/html/_static/scan_parameters/IPCAS_5_scantable.pdf

*Summary:* Images acquired on a 3-Tesla Siemens TrioTim scanner, 12-channel headcoil. 3D MPRAGE anatomic T1-weighted sequence (FA = 7°, TI = 1100 ms, TE = 3.5 ms, TR = 2530 ms, in-plane resolution = 1.0 × 1.0 $mm^2$, slice thickness = 1.0 mm, slice gap = 0.5 mm, FOV = 256 × 256 $mm^2$, number of slices = 176)
*Number of subjects included from this dataset:* 22

12. **Institute of Psychology, Chinese Academy of Sciences (IPCAS6)**
*Detailed acquisition parameters:*
http://fcon_1000.projects.nitrc.org/indi/CoRR/html/_static/scan_parameters/IPCAS_6_scantable.pdf
*Summary:* Images acquired on a 3-Tesla Siemens TrioTim scanner, 8-channel headcoil. 3D MPRAGE anatomic T1-weighted sequence (FA = 9°, TI = 900 ms, TE = 2.52 ms, TR = 1900 ms, in-plane resolution = 1.0 × 1.0 $mm^2$, slice thickness = 1.0 mm, slice gap = 0.5 mm, FOV = 256 × 246 $mm^2$, number of slices = 176)
*Number of subjects included from this dataset:* 2

13. **Institute of Psychology, Chinese Academy of Sciences (IPCAS8)**
*Detailed acquisition parameters:*
http://fcon_1000.projects.nitrc.org/indi/CoRR/html/_static/scan_parameters/IPCAS_8_scantable.pdf
*Summary:* Images acquired on a 3-Tesla Siemens TrioTim scanner, 12-channel headcoil. 3D MPRAGE anatomic T1-weighted sequence (FA = 7°, TI = 1100 ms, TE = 3.39 ms, TR = 2530 ms, in-plane resolution = 1.3 × 1.0 $mm^2$, slice thickness = 1.3 mm, slice gap = 0.65 mm, FOV = 256 × 256 $mm^2$, number of slices = 128)
*Number of subjects included from this dataset:* 13

14. **Jinling Hospital, Nanjing University (JHNU)**
*Detailed acquisition parameters:*
http://fcon_1000.projects.nitrc.org/indi/CoRR/html/_static/scan_parameters/JHNU_1_scantable.pdf
*Summary:* Images acquired on a 3-Tesla Siemens TrioTim scanner, 8-channel headcoil. 3D MPRAGE anatomic T1-weighted sequence (FA = 9°, TI = 900 ms, TE = 2.98 ms, TR = 2300 ms, in-plane resolution = 1.0 × 1.0 $mm^2$, slice thickness = 1.0 mm, slice gap = 0 mm, FOV = 256 × 256 $mm^2$, number of slices = 176)
*Number of subjects included from this dataset:* 30

15. **Ludwig-Maximilians-University (LMU1)**
*Detailed acquisition parameters:*
http://fcon_1000.projects.nitrc.org/indi/CoRR/html/_static/scan_parameters/LMU_1_scantable.pdf
*Summary:* Images acquired on a 3-Tesla Philips Achieva scanner, 32-channel headcoil. 3D T1-TFE anatomic T1-weighted sequence (FA = 8°, TI = 900 ms, TE = - ms, TR = 2375 ms, in-plane resolution = 1.0 × 1.0 $mm^2$, slice thickness = 1.0 mm, slice gap = 0 mm, FOV = 240 × 187 $mm^2$, number of slices = 220)
*Number of subjects included from this dataset:* 27

16. **Ludwig-Maximilians-University (LMU2)**
*Detailed acquisition parameters:*
http://fcon_1000.projects.nitrc.org/indi/CoRR/html/_static/scan_parameters/LMU_2_scantable.pdf
*Summary:* Images acquired on a 3-Tesla Siemens Verio scanner, 12-channel headcoil. 3D MPRAGE anatomic T1-weighted sequence (FA = 9°, TI = 900 ms, TE = 3.06 ms, TR = 2400 ms, in-plane resolution = 1.0 × 1.0 $mm^2$, slice thickness = 1.0 mm, slice gap = 0.5 mm, FOV = 256 × 246 $mm^2$, number of slices = 160)
*Number of subjects included from this dataset:* 33

17. **Ludwig-Maximilians-University (LMU3)**
*Detailed acquisition parameters:*
http://fcon_1000.projects.nitrc.org/indi/CoRR/html/_static/scan_parameters/LMU_3_scantable.pdf
*Summary:* Images acquired on a 3-Tesla Siemens TrioTim scanner, 12-channel headcoil. 3D MPRAGE anatomic T1-weighted sequence (FA = 9°, TI = 900 ms, TE = 3.06 ms, TR = 2400 ms, in-plane resolution = 1.0 × 1.0 $mm^2$, slice thickness = 1.0 mm, slice gap = 0.5 mm, FOV = 256 × 246 $mm^2$, number of slices = 256)
*Number of subjects included from this dataset:* 24

18. **Mind Research Network (MRN)**
*Detailed acquisition parameters:*
http://fcon_1000.projects.nitrc.org/indi/CoRR/html/_static/scan_parameters/MRN_1_scantable.pdf

*Summary:* Images acquired on a 3-Tesla Siemens TrioTrim scanner, 12-channel headcoil. MEMPR anatomic T1-weighted sequence (FA = 7°, TI = 1200 ms, TE = 1.64/3.5/5.36/7.22/9.08 ms, TR = 2530 ms, in-plane resolution = 1.0 × 1.0 $mm^2$, slice thickness = 1.0 mm, slice gap = 0.5 mm, FOV = 256 × 256 $mm^2$, number of slices = 192)
*Number of subjects included from this dataset:* 52

19. **Nathan Kline Institute (NKI1)**
*Detailed acquisition parameters:*
http://fcon_1000.projects.nitrc.org/indi/enhanced/NKI_MPRAGE.pdf
*Summary:* Images acquired on a 3-Tesla Siemens TrioTim scanner. 3D MPRAGE anatomic T1-weighted sequence (FA = 9°, TI = 900 ms, TE = 2.52 ms, TR = 1900 ms, in-plane resolution = 1.0 × 1.0 $mm^2$, slice thickness = 1.0 mm, slice gap = 0 mm, FOV = 250 × 250 $mm^2$, number of slices = 176)
*Number of subjects included from this dataset:* 23

20. **New York University (NYU1)**
*Detailed acquisition parameters:*
http://fcon_1000.projects.nitrc.org/indi/CoRR/html/_static/scan_parameters/nyu/nyu_anat.pdf
*Summary:* Images acquired on a 3-Tesla Siemens Magnetom Allegra scanner. Anatomic T1-weighted sequence (FA = 7°, TI = 1100 ms, TE = 3.25 ms, TR = 2530 ms, in-plane resolution = 1.3 × 1.0 $mm^2$, slice thickness = 1.3 mm, slice gap = 0 mm, FOV = 256 × 256 $mm^2$, number of slices = 128)
*Number of subjects included from this dataset:* 23

21. **Southwest University (SWU1)**
*Detailed acquisition parameters:*
NA
*Summary:* Anatomic T1-weighted images (FA = 9°, TI = 900 ms, TE = 2.52 ms, TR = 1900 ms, in-plane resolution = 1.0 × 0.97 $mm^2$, slice thickness = 0.97 mm, slice gap = NA, FOV = 176 × 250 $mm^2$, number of slices = 256)
*Number of subjects included from this dataset:* 17

22. **Southwest University (SWU2)**
*Detailed acquisition parameters:*
NA
*Summary:* Anatomic T1-weighted images (FA = 9°, TI = 900 ms, TE = 2.52 ms, TR = 1900 ms, in-plane resolution = 1.0 × 0.97 $mm^2$, slice thickness = 0.97 mm, slice gap = NA, FOV = 176 × 250 $mm^2$, number of slices = 256)
*Number of subjects included from this dataset:* 23

23. **Southwest University (SWU3)**
*Detailed acquisition parameters:*
NA
*Summary:* Anatomic T1-weighted images (FA = 9°, TI = 900 ms, TE = 2.52 ms, TR = 1900 ms, in-plane resolution = 1.0 × 0.97 $mm^2$, slice thickness = 0.97 mm, slice gap = NA, FOV = 176 × 250 $mm^2$, number of slices = 256)
*Number of subjects included from this dataset:* 20

24. **Southwest University (SWU4)**
*Detailed acquisition parameters:*
NA
*Summary:* Anatomic T1-weighted images (FA = 9°, TI = 900 ms, TE = 2.52 ms, TR = 1900 ms, in-plane resolution = 1.0 × 1.0 $mm^2$, slice thickness = 1.0 mm, slice gap = NA, FOV = 176 × 256 $mm^2$, number of slices = 256)
*Number of subjects included from this dataset:* 172

25. **University of McGill (UM)**
*Detailed acquisition parameters:*
NA
*Summary:* Anatomic T1-weighted sequence (FA = 9°, TI = 900 ms, TE = 2.98 ms, TR = 2300 ms, in-plane resolution = 1.0 × 1.0 $mm^2$, slice thickness = 1.0 mm, slice gap = NA, FOV = 176 × 240 $mm^2$, number of slices = 256)
*Number of subjects included from this dataset:* 78

26. **University of Pittsburg School of Medicine (UPSM)**

*Detailed acquisition parameters:*
http://fcon_1000.projects.nitrc.org/indi/CoRR/html/_static/scan_parameters/UPSM_1_scantable.pdf
*Summary:* Images acquired on a 3-Tesla Siemens TrioTim scanner, 12-channel headcoil. 3D MPRAGE anatomic T1-weighted sequence (FA = 8°, TI = 1050 ms, TE = 3.43 ms, TR = 2100 ms, in-plane resolution = 1.0 × 1.0 $mm^2$, slice thickness = 1.0 mm, slice gap = 0.5 mm, FOV = 256 × 256 $mm^2$, number of slices = 192)
*Number of subjects included from this dataset:* 89

27. **University of Utah (Utah1)**
*Detailed acquisition parameters:*
http://fcon_1000.projects.nitrc.org/indi/CoRR/html/_static/scan_parameters/utah/utah_anat.pdf
*Summary:* Images acquired on a 3-Tesla Siemens TrioTim scanner. 3D MPRAGE anatomic T1-weighted sequence (FA = 9°, TI = 900 ms, TE = 2.91 ms, TR = 2300 ms, in-plane resolution = 1.0 × 1.0 $mm^2$, slice thickness = 1.2 mm, FOV = 256 × 256 $mm^2$, number of slices = 160)
*Number of subjects included from this dataset:* 22

28. **University of Wisconsin, Madison (UWM)**
*Detailed acquisition parameters:*
NA
*Summary:* Anatomic T1-weighted sequence (FA = 12°, TI = 450 ms, TE = 3.18 ms, TR = 8.13 ms, in-plane resolution = 1.0 × 1.0 $mm^2$, slice thickness = 1.0 mm, slice gap = NA, FOV = 256 × 256 $mm^2$, number of slices = 156)
*Number of subjects included from this dataset:* 16

29. **Xuanwu Hospital, Capital University of Medical Sciences (XHCUMS)**
*Detailed acquisition parameters:*
http://fcon_1000.projects.nitrc.org/indi/CoRR/html/_static/scan_parameters/beijing_li/beijing_li_all.pdf
*Summary:* Images acquired on a 3-Tesla Siemens TrioTim scanner. 3D MPRAGE anatomic T1-weighted sequence (FA = 9°, TI = 800 ms, TE = 2.15 ms, TR = 1600 ms, in-plane resolution = 1.0 × 1.0 $mm^2$, slice thickness = 1.0 mm, slice gap = 0 mm, FOV = 256 × 256 $mm^2$, number of slices = 176)
*Number of subjects included from this dataset:* 17

*Summary of final inclusions from CoRR:*

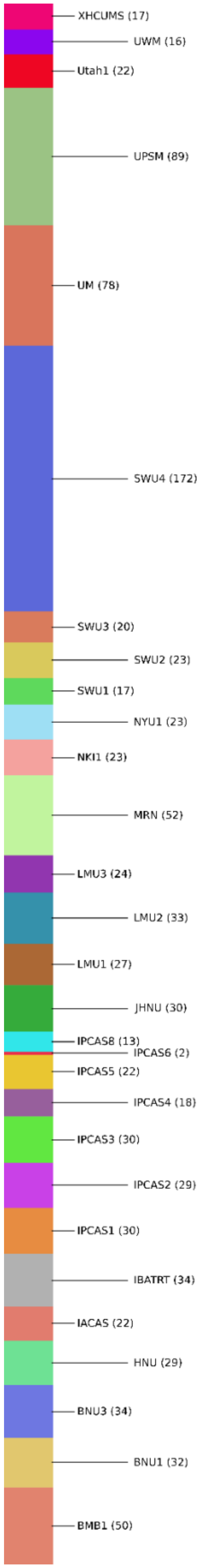

#### Dallas Lifespan Brain Study (DLBS)

**Final number of healthy subjects included in the study: 269**

*Website:* http://fcon_1000.projects.nitrc.org/indi/retro/dlbs.html

*Description:* The Dallas Lifespan Brain Study (DLBS) is a major effort designed to understand the antecedents of preservation and decline of cognitive function at different stages of the adult lifespan, with a particular interest in the early stages of a healthy brain's march towards Alzheimer Disease. The DLBS aims to (1) understand how brain atrophy as well as accumulation of amyloid and tau affects healthy aging and cognition, (2) understand how the brain develops "neural scaffolds" to support cognition, and also (3) develop a corpus of research on the cognitive neuroscience of middle age. We have enrolled 350 healthy adults, aged 20-89 in the study who are thoroughly characterized in terms of cognition, brain structure and brain function across the adult lifespan. At Wave 1, DLBS participants received a structural MRI with DTI, three task-based functional MRI scans, and a resting state scan on a Philips 3T scanner and extensive cognitive testing combined with a detailed psychosocial battery. Amyloid imaging, using F-18 Florbetapir was conducted on a Siemens ECAT HR PET scanner on 60% of the sample during Wave 1. The DLBS is one of the most complete studies of the aging mind that is available, particularly in the United States, and can address many important hypotheses regarding the cognitive neuroscience of aging.

*Image acquisition:* Images acquired on a 3-Tesla Philips Achieva scanner, 8-channel headcoil. MPRAGE T1-weighted sequence (FA = 12°, TE = 3.7 ms, TR = 8.1 ms, in-plane resolution = 1 × 1 $mm^2$, slice thickness = 1 mm, slice gap = 0 mm, FOV = 204 × 256 $mm^2$, number of slices = 160)

#### Brain Genomics Superstruct Project (GSP)

**Final number of healthy subjects included in the study: 1552**

*Website:* https://www.neuroinfo.org/gsp/

*Description:* Large scale imaging data sets are necessary to address complex questions regarding the relationship between brain and behavior. The Brain Genomics Superstruct Project Open Access Data Release exposes a carefully vetted collection of neuroimaging, behavior, cognitive, and personality data for over 1,500 human participants. Each neuroimaging data set includes one high-resolution Magnetic Resonance Imaging (MRI) acquisition and one or more resting-state functional MRI acquisitions. Each functional acquisition is accompanied by a fully-automated quality assessment and pre-computed brain morphometrics.

*Image acquisition:* Images acquired on a 3-Tesla Siemens TrioTim scanner, 12-channel headcoil. MEMPRAGE T1-weighted sequence (FA = 7°, TR = 2200 ms, TI = 1100 ms, TE1 = 1.54 ms, TE2 = 3.36 ms, TE3 = 5.18 ms, TE4 = 7 ms, in-plane resolution = 1.2 × 1.2 $mm^2$, slice thickness = 1.2 mm, slice gap = 0 mm, FOV = 230 × 230 $mm^2$, number of slices = 144).

Further details available at:

https://static1.squarespace.com/static/5b58b6da7106992fb15f7d50/t/5b68650d8a922db3bb807a90/1533568270847/GSP_README_140630.pdf

#### Human Connectome Project (HCP)

**Final number of healthy subjects included in the study: 1113**

*Website:* https://www.humanconnectome.org

*Description:* Mapping the human brain is one of the great scientific challenges of the 21st century. The Human Connectome Project (HCP) has tackled key aspects of this challenge by charting the neural pathways that underlie brain function and behavior, including high-quality neuroimaging data in over 1100 healthy young adults. Using greatly improved methods for data acquisition, analysis, and sharing, the HCP has provided the scientific community with data and discoveries that greatly enhance our understanding of human brain structure, function, and connectivity and their relationships to behavior.

*Image acquisition:* Images acquired on a 3-Tesla Siemens Skyra custom scanner, 32-channel headcoil. MPRAGE T1-weighted sequence (FA = 8°, TR = 2400 ms, TE = 2.14 ms, TI = 1000 ms, in-plane resolution = 0.7 × 0.7 $mm^2$, slice thickness = 0.7 mm, slice gap = 0 mm, FOV = 224 × 224 $mm^2$, number of slices = 256, 2 averages). Further details can be found in (5-7).

#### Information eXtraction from Images (IXI)

**Final number of healthy subjects included in the study: 438**

*Website:* http://brain-development.org/ixi-dataset/

*Description:* In this project we have collected nearly 600 MR images from normal, healthy subjects. The MR image acquisition protocol for each subject includes:

- T1, T2 and PD-weighted images
- MRA images
- Diffusion-weighted images (15 directions)

The data has been collected at three different hospitals in London:

- Hammersmith Hospital using a Philips 3T system (HH) *(Number of subjects from this site: 130)*
- Guy's Hospital using a Philips 1.5T system (GH) *(Number of subjects from this site: 261)*
- Institute of Psychiatry using a GE 1.5T system (IOP) *(Number of subjects from this site: 47)*

*Image acquisition:* Hammersmith Hospital: images acquired on a 3-Tesla Philips Intera scanner. T1-weighted sequence (FA = 8°, TR = 9.6 ms, TE = 4.6 ms, in-plane resolution = 0.937 × 0.937 $mm^2$, slice thickness = 1.2 mm, slice gap = NA, FOV = 240 × 240 $mm^2$, number of slices = 150). Guy's Hospital: images acquired on a 1.5 Tesla Philips Gyroscan scanner. T1-weighted sequence (FA = 8°, TR = 9.8 ms, TE = 4.6 ms, in-plane resolution = 0.937 × 0.937 $mm^2$, slice thickness = 1.2 mm, slice gap = NA, FOV = 240 × 240 $mm^2$, number of slices = 150).

#### MPI-Leipzig Mind Brain Body (MPI-LMBB)

**Final number of healthy subjects included in the study: 292**

*Website:* https://openneuro.org/datasets/ds000221

*Description:* The MPI-Leipzig Mind-Brain-Body dataset contains MRI and behavioral data from 318 participants. Datasets for all participants include at least a structural quantitative T1-weighted image and a single 15-minute eyes-open resting-state fMRI session. The participants took part in one or two extended protocols: Leipzig Mind-Body-Brain Interactions (LEMON) and Neuroanatomy & Connectivity Protocol (N&C).

*Image acquisition:* Images acquired on a 3-Tesla Siemens Verio scanner, 32-channel headcoil. MP2RAGE T1-weighted sequence (FA1 = 4°, FA2 = 5°, TR = 5000 ms, TE = 2.92 ms, TI1 = 700 ms, TI2 = 2500 ms, in-plane resolution = 1 × 1 $mm^2$, slice thickness = 1 mm, slice gap = 0 mm, FOV = 256 × 240 $mm^2$, number of slices = 176). Further details in (8).

#### Enhanced Nathan Kline Institute-Rockland Sample (NKI-RS)

**Final number of healthy subjects included in the study: 350**

*Website:* http://fcon_1000.projects.nitrc.org/indi/enhanced/

*Description*: The enhanced Nathan Kline Institute-Rockland Sample (NKI-RS) is an ongoing, institutionally centered endeavor aimed at creating a large-scale (N > 1000) community sample of participants across the lifespan. Measures include a wide array of physiological and psychological assessments, genetic information, and advanced neuroimaging. Anonymized data will be publicly shared openly and prospectively (i.e., on a quarterly basis). The National Institute of Mental Health strategic plan for advancing psychiatric neuroscience calls for increased discovery science initiatives in an effort to deliniate developmental trajectories for risk and resilience across the lifespan. Such initiatives require large, openly accessible datasets that contain sufficient detail to enable the generation and testing of multiple hypotheses. The NKI-RS is intended to help meet this need.

*Image acquisition:* Scans part of the cross-sectional lifespan connectomics and longitudinal developmental connectomics protocol were acquired on a 3-Tesla Siemens Magnetom TrioTim scanner. MPRAGE T1-weighted sequence (FA = 9°, TR = 1900 ms, TE = 2.52 ms, TI = 900 ms, in-plane resolution = 1 × 1 $mm^2$, slice thickness = 1 mm, slice gap = 0 mm, FOV = 250 × 250 $mm^2$, number of slices = 176). Scans part of the Real-time neurofeedback protocol were acquired on a 3-Tesla Siemens Magnetom TrioTim scanner. MPRAGE T1-weighted sequence (FA = 8°, TR = 2600 ms, TE = 3.02 ms, TI = 900 ms, in-plane resolution = 1 × 1 $mm^2$, slice thickness = 1 mm, slice gap = 0 mm, FOV = 256 × 256 $mm^2$, number of slices = 192). Further details in (9).

#### Southwest University Adult Lifespan Dataset (SALD)

**Final number of healthy subjects included in the study: 415**

*Website:* http://fcon_1000.projects.nitrc.org/indi/retro/sald.html

*Description*: Some of the most urgent issues confronting the scientific community today involve mental health and the development of the human brain. Thus, the field of developmental neuroscience aims to uncover the developmental trajectory of the human brain and understand the changes that occur as a function of aging. Understanding the mechanisms of aging will help the scientific community move closer towards discovering the causes of nervous system diseases related to aging. Here, we describe the data generated in the Southwest University Adult Lifespan Dataset (SALD), which comprises a large cross-sectional sample (n = 494; age range = 19-80) undergoing a multi-modal (sMRI, rs-fMRI, and behavioral). The goals of the SALD are to give researchers the opportunity to map the structural and functional changes the human brain undergoes throughout adulthood and to replicate previous findings.

*Image acquisition:* Images were acquired on a 3-Tesla Siemens Magnetom TrioTim scanner. MPRAGE T1-weighted sequence (FA = 9°, TR = 1900 ms, TE = 2.52 ms, TI = 900 ms, in-plane resolution = 1 × 1 $mm^2$, slice thickness = 1 mm, slice gap = 0 mm, FOV = 256 × 256 $mm^2$, number of slices = 176). Further details in (10).

### Data preparation

The overall data preparation process is illustrated in Supplemental Figure 1. This included the following processing and quality assessment steps.

#### Image processing

T1 images were processed with Freesurfer (v6.0, https://surfer.nmr.mgh.harvard.edu) using Northwestern University's High Performance Computing Cluster (QUEST, https://www.it.northwestern.edu/research/user-services/quest/) and the CBRAIN platform (11). Preprocessing steps included bias field correction, intensity normalization, spatial normalization, skull stripping and tissue segmentation (12). The inner cortical surface, matching the white matter / grey matter junction, and the outer cortical surface, matching the grey matter / cerebro-spinal fluid interface, were then extracted. The surfaces were corrected for possible topological defects, inflated and parameterized (13, 14).

#### Quality assessment

Subjects were excluded from analysis if Freesurfer could not complete surface extraction. No attempt was made to reprocess subjects whose Freesurfer process failed before completion. For completed Freesurfer data, an automated process mapped the cortical thickness on the inner and outer cortical surfaces, generated a lateral and a medial view of each surface, and concatenated all views in a single image. These images were visually inspected to ensure that the surfaces contained no obvious large errors. If a significant error was suspected, the original T1 image and four extracted cortical surfaces were opened simultaneously using freeview to make the final decision on the quality.

Subjects were excluded if at least one large error, significant enough to globally modify the sulcal pattern was identified. For example, subjects were discarded if one or both temporal poles were not entirely extracted, widespread inclusion of dura or if the surface was particularly noisy. Localized errors such as the inclusion of dura at the crown of a gyrus or local roughness were not disqualifying.

Finally, affine registrations to MNI space were inspected for all subjects. This was done by visually reviewing automated snapshots of the cortical surfaces aligned with the MNI space. Incorrect registrations were corrected by re-registering individual's skull stripped native image to skull stripped MNI305 template. New registrations were again visually checked and data were included if they were correct.

##### Registration of the cortical surfaces to the MNI space

The four extracted cortical surfaces (left and right hemispheres, inner and outer cortical surfaces) were normalized by registering them to the MNI space (MNI305 template). This was done by applying the affine registration to the surface vertices coordinates in order to spatially normalize the cortical surfaces and normalize brain volumes.

#### Surface registration

The purpose of surface registration is to establish a correspondence across individuals. By default, Freesurfer performs a non-rigid surface registration to improve cortical folding alignment (as determined by the curvature of the surfaces) between a subject and the surface template (14). This can result in distortion of the sulcal pattern in order to better match the surface template. To overcome this and preserve the initial sulcal folding pattern, surfaces were rigidly registered to the surface template (Freesurfer's *fsaverage*). In a trade-off between precision of cortical alignment across subjects and preservation of individual subjects' cortical folding pattern, this maneuver prioritized preservation of subjects' folding pattern.

#### Resampling surface coordinates to the common surface

Cartesian coordinates of the vertices in individual's native space were registered to the MNI space by applying the affine transformation, then they were registered to the common surface space by applying the rigid surface transformation. Finally, vertices coordinates were resampled to the freesurfer's *fsaverage5* template to decrease the number of vertices in the common surface space. This resulted in 10,242 vertices per cortical surface, with a one-to-one correspondence between the inner and outer cortical surfaces in each hemisphere.

### Network and Training Parameters

#### Sex prediction

As a first experiment, the network was trained to classify sex. Training and validation were handled via a $k$-fold cross-validation approach with $k$ = 5, therefore the network was trained with 5128 instances and validated with 1282 independent instances. For this task, the loss function was cross-entropy, $F_C$ was set to 32 filters, the number of blocks $B$ set to 4 and $F_L$ was set to 128. In addition, the polynomial order was set to 5 for all filters, the learning rate was set to 0.001 with an exponential decay 0.95 every 400 iterations, filter weights were $L2$ regularized to 5 × $10^{-4}$, dropout 0.5 was applied to the fully connected layer, the batch size was 64 and the optimization was performed with ADAM procedure (15).

#### Age prediction

As a second task, the network was trained for age prediction using the same $k$-fold approach. The loss function was the mean squared error, $F_C$ was set to 32 filters, the number of blocks $B$ was 7, $F_L$ was 16 and the polynomial order was 4 for all filters. The initial learning rate was 0.001 with an exponential decay 0.98 every 400 iterations, filter weights $L2$ regularization set to 1 × $10^{-5}$, dropout 0.5 was applied to the FC layer, the batch size was set to 64 and optimization was done with the ADAM method.

### References


1. Karayanidis F*, et al.* (2016) The Age-ility Project (Phase 1): Structural and functional imaging and electrophysiological data repository. *NeuroImage* 124:1137-1142.
2. Taylor JR*, et al.* (2017) The Cambridge Centre for Ageing and Neuroscience (Cam-CAN) data repository: Structural and functional MRI, MEG, and cognitive data from a cross-sectional adult lifespan sample. *NeuroImage* 144:262-269.
3. Shafto MA*, et al.* (2014) The Cambridge Centre for Ageing and Neuroscience (Cam-CAN) study protocol: a cross-sectional, lifespan, multidisciplinary examination of healthy cognitive ageing. *BMC Neurology* 14:204.
4. Zuo X-N*, et al.* (2014) An open science resource for establishing reliability and reproducibility in functional connectomics. *Scientific Data* 1:140049.
5. Glasser MF*, et al.* (2013) The minimal preprocessing pipelines for the Human Connectome Project. *Neuroimage* 80:105-124.
6. Ugurbil K*, et al.* (2013) Pushing spatial and temporal resolution for functional and diffusion MRI in the Human Connectome Project. *Neuroimage* 80:80-104.
7. Sotiropoulos SN*, et al.* (2013) Advances in diffusion MRI acquisition and processing in the Human Connectome Project. *NeuroImage* 80(0):125-143.
8. Mendes N*, et al.* (2017) A functional connectome phenotyping dataset including cognitive state and personality measures.
9. Nooner K*, et al.* (2012) The NKI-Rockland Sample: A Model for Accelerating the Pace of Discovery Science in Psychiatry. 6(152).
10. Wei D*, et al.* (2018) Structural and functional MRI from a cross-sectional Southwest University Adult lifespan Dataset (SALD). *bioRxiv*.
11. Sherif T*, et al.* (2014) CBRAIN: a web-based, distributed computing platform for collaborative neuroimaging research. 8(54).
12. Dale AM, Fischl B, & Sereno MI (1999) Cortical Surface-Based Analysis: I. Segmentation and Surface Reconstruction. *NeuroImage* 9(2):179-194.
13. Fischl B, Sereno MI, & Dale AM (1999) Cortical Surface-Based Analysis: II: Inflation, Flattening, and a Surface-Based Coordinate System. *NeuroImage* 9(2):195-207.
14. Fischl B, Sereno MI, Tootell RB, & Dale AM (1999) High-resolution intersubject averaging and a coordinate system for the cortical surface. *Human brain mapping* 8(4):272-284.
15. Kingma DP & Ba JJapa (2014) Adam: A method for stochastic optimization.

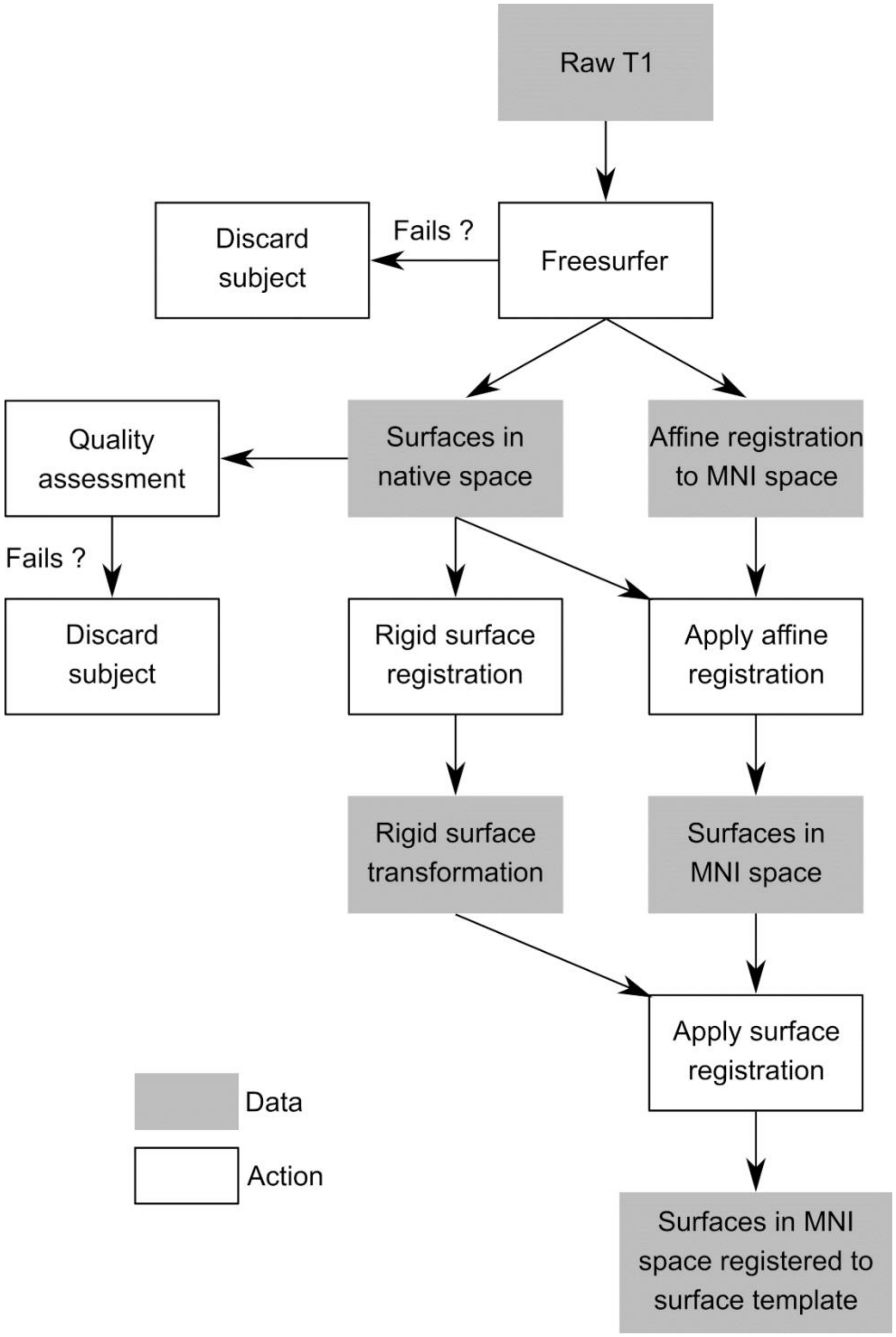


**Supplemental Figure 1.** Flowchart of data preparation. After cortical surfaces are extracted using Freesurfer, they are affinely registered to the MNI space and rigidly registered with the surface template to ensure vertex-wise correspondence. Quality assessment ensures that the quality of surface extraction and registration is satisfactory.
