## Supplemental tables for "Geometric deep learning on brain shape predicts sex and age"

| **Dataset** | **Inner** | **Inner & outer** |
| --- | --- | --- |
| ABIDE-II | 0.83 | 0.81 |
| CamCan | 0.89 | 0.90 |
| CoRR | 0.81 | 0.83 |
| DLBS | 0.84 | 0.89 |
| GSP | 0.87 | 0.90 |
| HCP | 0.86 | 0.90 |
| IXI | 0.85 | 0.89 |
| MPI-LMBB | 0.86 | 0.87 |
| NKIRS | 0.85 | 0.88 |
| SALD | 0.76 | 0.80 |
| age-ility | 0.84 | 0.89 |

**Table 1.** Accuracy of sex classification per dataset.

| **Cohort** | **Inner** | **Inner & outer** |
| --- | --- | --- |
| ABIDE-II | 5.06 | 4.88 |
| CamCan | 7.03 | 6.21 |
| CoRR | 4.82 | 4.50 |
| DLBS | 8.61 | 7.94 |
| GSP | 2.86 | 2.73 |
| HCP | 3.86 | 3.65 |
| IXI | 6.61 | 6.24 |
| MPI-LMBB | 5.50 | 4.98 |
| NKIRS | 6.34 | 6.22 |
| SALD | 6.76 | 6.19 |
| age-ility | 3.87 | 3.64 |

**Table 2.** Mean absolute error for age prediction per dataset.
